# Bactericidal activity of ZnO nanoparticles-anti TB drugs combination towards H37Rv strain and multidrug-resistant isolates of *Mycobacterium tuberculosis* via SufB splicing inhibition

**DOI:** 10.1101/2025.03.13.643114

**Authors:** Deepak Kumar Ojha, Ashwaria Mehra, Sunil Swick Rout, Sidhartha Giri, Sasmita Nayak

## Abstract

Tuberculosis (TB) remains a significant global health threat, claiming millions of lives annually. Despite advancements in treatment, the emergence of drug-resistant strains has hindered effective TB control. The current management for TB is prolonged with severe side effects, leading to poor patient compliance. Metal-based nanoparticles are shown to manage drug-sensitive TB when combined with anti-TB drugs. However, mycobactericidal potential of nanoparticles towards drug-resistant TB is not confirmed yet. This work explores the bactericidal potential of Zinc Oxide Nanoparticles (ZnONPs, 40 nm) in managing both drug-sensitive and drug-resistant TB in combination with anti-TB drugs. It was found that ZnONPs inhibit generation of active SufB protein via splicing inhibition, an essential event for *Mycobacterium tuberculosis* (*Mtb*) survival. While TEM and UV-visible spectroscopy identified NPs∼protein interaction, SEM visualised extensive membrane damage in H37Rv and multidrug-resistant (MDR) *Mtb* cells. Alamar blue assay and spread plate method detected minimum inhibitory concentration and minimum bactericidal concentration of ZnONPs towards H37Rv strain and MDR *Mtb* isolates. *In vitro* studies identified a combination with ZnONPs that reduced effective doses for anti-TB drugs towards H37Rv and MDR *Mtb* isolates.

A correlation to splicing inhibition was made by performing Alamar blue assay in SufB intein-less microbe, *Mycobacterium smegmatis*. A similar drug combination, attenuated the mycobacterial load, inflammation in the spleen & lungs, and protected against *Mtb* induced splenomegaly in infected mice. Thus, ZnONPs can be used as potent additive in anti-TB regimen to manage drug-susceptible and drug-resistant TB, addressing challenges such as prolonged therapy, drug toxicity and poor patient compliance.

## 1. Introduction

Tuberculosis (TB) is one of the leading causes of death worldwide, ranking among the top ten deadliest diseases globally, and is the single most lethal infectious agent, surpassing even HIV/AIDS ^1,2^. However, efforts to combat TB are often impeded by several challenges, including absence of an effective vaccine, the complexities of treatment regimens, and the rapid emergence of drug-resistant *Mycobacterium tuberculosis* (*Mtb*) strains ^3,4^. Moreover, prolonged therapy can lead to the phenomenon of cross-resistance, where bacteria become resistant to multiple drugs by exploiting shared mechanisms of action ^5^. The standard treatment for drug-susceptible and drug-resistant TB spans approximately 6 months and over 18 months, respectively. The intricacy of these regimens, along with drug-induced side effects, often leads to poor patient compliance, worsening the current situation due to the emergence of new drug-resistant strains ^6^. Hence, there is an urgency to develop next-generation strategies to mitigate the limitations and enhance the accessibility of TB management.

One promising approach to control TB could be targeting the intein splicing, which is a critical intracellular process that enables *Mycobacterium tuberculosis* (*Mtb*) survival and persistence by generating essential proteins ^7,8^. Inteins are intervening polypeptides co-translated within the host precursor proteins and self-excised through protein splicing to generate active protein^9^. Given, that several human pathogens harbour inteins within vital proteins, and intein splicing is critical for their survival, the regulation of splicing can function as a promising target for developing anti-microbial drugs especially towards drug-resistant organisms. *Mtb* SufB, which is an Fe-S cluster assembly protein represents a notable example among mycobacterial proteins, for its significance in disease pathogenesis ^10,11^. The SUF system as a whole is required for metalloprotein biogenesis and plays an important role in mycobacterial survival during iron limitation including oxidative and nitrosative stresses within infected macrophages^12,13^.

In recent years, the synergistic integration of nanotechnology and medicine has emerged as a popular and diverse approach, particularly for the development of anti-microbial compounds. The convergence of these disciplines has demonstrated significant potential in addressing various challenges associated with bacterial infections ^14–16^. Further, the interaction between nanoparticles and proteins holds significant importance in various areas of contemporary biomedical research, particularly in the fields of nanomedicine and nano diagnostics ^17–21^. When nanoparticles interact with proteins, they form a protein corona, which alters the biological activity of the bound protein. This phenomenon has profound implications for understanding the behaviour of nanoparticles within biological environments and their potential applications in drug delivery, diagnostics, and therapeutics ^22–29^.

Thus, nanotechnology is gaining popularity as the cornerstone of modern research and therapeutics. This frequently involves metal-based NPs like ZnONPs that offer varied uses in emerging applications such as antimicrobial, anti-inflammatory, anti-tumour, and in wound healing ^30–35^. ZnONPs exhibit superior bioactivity, primarily due to their small size, and increased surface area to volume ratio ^36^. ZnONPs are also preferred as efficient drug delivery vehicles for various diseases including cancer, because of their safety, stability, and low cost^36,37^.

Previous research have shown the bacteriostatic effects of ZnONPs towards both drug-sensitive and extensively drug-resistant (XDR) *Mtb* strains at concentrations equal to or higher than 1 µg/ mL ^38^. The same study has also determined 4 µg/mL as the inhibitory concentration of ZnONPs against the multidrug-resistant (MDR) strain of *Mtb*. Another work has reported the bactericidal role of ZnONPs - Se against *Mtb* BCG and H37Rv strains ^39^. The antimicrobial mechanisms for ZnONPs effects are mostly attributed to a cumulative effect of cell membrane disruption, ROS production, release of toxic Zn^2+^ ions, and toxicity induced by their photocatalytic action ^40^. Although ZnONPs are known to exhibit potent antimicrobial and fungicidal activity towards different microbes ^41^, their bactericidal potential on drug-resistant *Mtb* strains are not shown till date. In a different context, inhibitory effects of divalent metal ions such as Zn^2+^, and Cu^2+^ on the splicing of various intein containing precursor proteins were explored extensively, but studies showing intein splicing attenuation in presence of metal-based NPs is missing till date ^8,42–47^.

Current study addresses the following questions: 1) Whether ZnONPs can regulate *Mycobacterium tuberculosis (Mtb)* SufB intein splicing and N-terminal cleavage reactions? 2) Since intein splicing generates a functional SufB protein that is critical for mycobacterial survival, whether ZnONPs mediated regulation of SufB splicing can affect the viability of whole cell *Mtb?* 3) Importantly, if, the viability of both drug-sensitive and drug-resistant strains of *Mtb* are influenced by ZnONPs? Next, 4) what is the minimum effective concentration of ZnONPs for the observed effect? 5) Whether ZnONPs combination can reduce the required dosage of existing anti-TB drugs to control both drug-sensitive and drug-resistant *Mtb* infection? 6) Whether ZnONPs exhibit a bacteriostatic or bactericidal effect towards drug-sensitive and drug-resistant *Mtb* strains*?* 7) Can ZnONPs protect against the effects of *Mtb* infection in an animal model?

In this work, ZnONPs (40 nm) activity on the splicing and N-terminal cleavage reactions of the *Mycobacterium tuberculosis* (*Mtb*) SufB precursor protein was examined. Initial cytotoxicity evaluation via MTT assay indicated that ZnONPs are biocompatible towards HEK293T cells (human embryonic kidney cell lines), up to a concentration of 50 µg/ml, consequently rest of the experiments were performed within this limit. UV-Vis spectroscopy confirmed ZnONPs ∼ SufB interaction, further visualised by Transmission Electron Microscopy (TEM), which identified protein “corona” formation around the ZnONPs. Additional experiments like Dynamic Light Scattering (DLS), and zeta potential confirmed ZnONPs-*Mtb* SufB interaction. Next, ZnONPs over varied concentration range (0.5 µg/ml-50 µg/ml) significantly inhibited the splicing and N-terminal cleavage reactions of *Mtb* SufB precursor protein as shown by *in vitro* protein refolding assay. SDS-PAGE and Western blot analyses visualized and confirmed the splicing and cleavage products. Further, Alamar blue assay and spread plate method detected anti-mycobacterial activity of ZnONPs with a minimum inhibitory concentration (MIC) / minimum bactericidal concentration (MBC) of 5 µg/ml/17 µg/ml and 14 µg/ml/23 µg/ml towards H37Rv strain (drug-sensitive) and multidrug-resistant (MDR) *Mtb* isolates respectively. In a comparative study, we also observed a reduction in the effective dose of anti-TB drugs INH/RIF and LFX/MXF in combination with ZnONPs that could efficiently kill H37Rv strain and multidrug-resistant (MDR) *Mtb* isolates respectively. The observed activity was correlated to splicing inhibition by performing Alamar blue assay in SufB intein-less microbe *Mycobacterium smegmatis*. Scanning Electron Microscopy (SEM) demonstrated mycobactericidal activity of ZnONPs towards both drug-sensitive and drug-resistant *Mtb* (H37Rv) strains. Finally, infected mice model study revealed that ZnONPs in combination with anti-TB drugs can effectively reduce mycobacterial load in the lungs and spleen. The observed effects were noticed when ZnONPs was administered as a solo therapy or in combination with a reduced dose of RIF compared to previously recommended amount of RIF. Likewise, the same combination of ZnONPs - RIF, not only attenuated the inflammation in lungs and spleen, but also provided resistance to splenomegaly in the infected mice. Therefore, we suggest the use of ZnONPs as a promising adjunctive therapy along with standard anti-TB drugs to combat both drug-sensitive and multidrug-resistant *Mtb* infections, by inhibiting the splicing of an essential protein, *Mtb* SufB. This approach may not only mitigate the toxicity associated with conventional and contemporary anti-TB regimens but also potentially reduce the duration of therapy, thereby enhancing patient compliance.

## 2. Materials and Methods

### 2.1 ZnO nanoparticles

ZnONPs (≥ 40 nm) were procured from Sigma Aldrich (Product #721077).

### 2.2 MTT Assay ^48–50^

The cytotoxic effects of ZnONPs were evaluated using the MTT (3-[4,5-dimethylthiazol-2-yl]-2,5-diphenyltetrazolium bromide) assay over a period of 72h. HEK239T cells (10^4^ cells/well) [Product # HEK239T (Human embryonic kidney cell line, ATCC (CRL-3216)] were seeded in Dulbecco’s Modified Eagle Medium (DMEM, Gibco, Invitrogen-2562497) in a 96-well plate (200 μl per well) and allowed to adhere for 24 hours at 37°C in 5% CO2 incubator. Subsequently, varying concentrations of ZnONPs (5 µg/ml, 14 µg/ml, 17 µg/ml, 23 µg/ml µg/ml, 26 µg/ml, 50 µg/ml, 100 µg/ml, 150 µg/ml) were added, followed by incubation at 37°C in 5% CO2 incubator for 72 hours. HEK239T cells (10^4^ cells/well) incubated in absence of nanoparticles was considered as the control for 100% viable cell growth. DMEM culture medium alone was used as blank control. 20 μl of MTT (EZcount^TM^ MTT cell assay kit, Himedia-CCK003) solution (5 mg ml^−1^ in incomplete medium) was added to each well and incubated for 3h to facilitate formation of purple formazan crystals under standard culture conditions. Next, MTT solution was discarded, and the crystals were solubilized by gently stirring on orbital shaker for 30 minutes. After solubilization, the absorbance (570 nm) of each well was measured by using the ELISA plate reader (Bio Teck Epoch Microplate Spectrophotometer). The percentage of cell viability was calculated using the following formula.

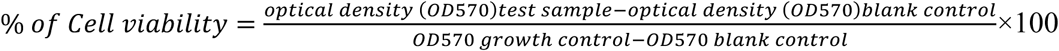

### 2.3 UV Visible spectroscopy ^51–53^

Purified *Mtb* SufB precursor protein was refolded in presence of different concentrations of ZnONPs (0.5 μg/ml, 26 μg/ml, and 50 μg/ml) as explained in section 2.7 for *in vitro* refolding assay. Then, the treated samples were analyzed via UV-Visible spectroscopy (UV-1800 Shimadzu) over the visible region of the spectrum (200-800 nm). To account for potential buffer-nanoparticle interactions, control samples included refolding buffer (20 mM sodium phosphate, 0.5 M NaCl, 0.5 M arginine, and 8 M urea) alone and the buffer with 50 µg/ml of nanoparticles incubated under the same conditions. The nanoparticle-treated buffer served as a blank for baseline correction. The absorbance results were analyzed by plotting absorbance versus wavelength using OriginPro 8.5 (Version 8.5, OriginLab Corporation, Northampton, MA, U.S.A.).

### 2.4 Quantification of NPs-bound proteins via Bradford assay ^54–56^

The quantification of adsorbed protein was conducted using the Bradford assay. *Mtb* SufB precursor protein was renatured *in vitro* both in presence and absence various concentrations of ZnONPs (0.5 µg/ml, 26 µg/ml, and 50 µg/ml) for 4 hours, as mentioned in section 2.7. Next, the protein-NPs mixture was centrifuged at 13,000 rpm for 30 minutes, at 4°C, to separate the protein-bound nanoparticles. Then, the pellet containing the protein ∼ NPs was resuspended in 100 µl of 1*×*PBS and analysed using Bradford reagent to quantify the amount of adsorbed proteins. Control samples included nanoparticles in 1*×*PBS and test proteins resuspended in 1*×*PBS.

### 2.5 TEM Analysis ^57,58^

*Mtb* SufB precursor protein-ZnONPs interaction was analysed by Transmission Electron Microscope (Jeol JEM-1400) for visualising protein corona formation. *Mtb* SufB precursor protein was overexpressed, purified and refolded with 50 µg/ml of ZnONPs as mentioned in section 2.7. The protein-ZnONPs complex was collected via centrifugation (13,000 rpm for 30 minutes) at 4^0^C, washed twice with phosphate buffer saline (1×PBS) and the supernatant was discarded. Then the pellet (protein-ZnONPs complex) was resuspended and fixed with 100 µl of 2.5% glutaraldehyde in 1×PBS for 1 hr. Finally, 5 μl of solution (ZnONP-complex) was placed onto the carbon-coated copper grid and imaged under TEM with an acceleration voltage of 120 Kv.

### 2.6 Purification of overexpressed proteins

The details of the plasmids used in this study are provided elsewhere ^7^. The *Mtb* SufB wild type precursor, and *Mtb* SufB splicing inactive double mutant (C1A/N359A) (splicing negative control), all carrying an N-terminal 6X His tag, were overexpressed in BL21 (DE3) *E. coli* cells via IPTG (500 μM, sigma 367-93-1) induction. The cells were resuspended in lysis buffer (20mM sodium phosphate, 0.5M NaCl, pH 7.4) and lysed using a tip sonicator (Sonics vibra cell VCX-130) ^59^. The cell lysate was centrifuged at 16,500xg for 20 minutes at 4°C and the supernatant was discarded. The collected IB materials were washed thrice via centrifugation in lysis buffer, solubilized in 8 M urea (Merck, 1084870500) buffer (lysis buffer, 8M urea, 10 mM imidazole (MP–biochemicals-288-32-4), and centrifuged at 16,500xg for 20 minutes at 4°C to collect the supernatant. The solubilized proteins were purified using a Ni-NTA affinity column (Ni-NTA His trap, HP GE healthcare life sciences-17524802) ^60–64^. Before sample application, the columns were equilibrated with binding buffer (20 mM sodium phosphate, 0.5 M NaCl, 8 M urea, 10 mM imidazole), and after loading the samples, the columns were washed several times (15 column volumes) with the wash buffer (20 mM sodium phosphate, 0.5M NaCl, 8M urea, 40 mM imidazole). Finally, the test proteins were eluted in elution buffer (20mM sodium phosphate, 0.5M NaCl, 500 mM imidazole, 8 M Urea, pH 7.4) and quantified using Bradford’s assay.

### 2.7 In vitro protein refolding and splicing analysis via SDS gradient PAGE

Denatured test proteins (2.5 µM) in 8M urea buffer were renatured in a refolding buffer containing 20 mM sodium phosphate, 0.5 M NaCl, 0.5 M L-Arginine (A5006 Sigma Aldrich), and 2 mM TCEP-HCl (Sigma-51805-45-9). Protein refolding was performed for 4 hours at 25°C ^59–64^ in the presence and absence of ZnO nanoparticles (ZnONPs) over a varied concentration range (0.5 µg/ml to 50 µg/ml). For the 0-hour sample, protein sample was collected before initiation of the refolding, followed by immediate addition of the loading dye (0.1% bromophenol blue, 50% glycerol, β-mercaptoethanol, 10% SDS, tris, pH 6.8) and rapid freezing at -20°C. After 4h of protein renaturation, rest of the samples were collected, loading dye was added, samples were boiled at 95°C for 5 minutes. Then protein products were resolved through 5∼10% gradient SDS-PAGE gel. The protein bands were visualized by staining with Coomassie brilliant blue R-250 (Catalogue no #112553, Sigma-Aldrich), and densitometric analysis was performed using the GelQuant.Net biochemical solutions software. The splicing and cleavage efficiencies were calculated as the percentage ratio of the total splicing product (LE and I) over the total proteins (LE+I+P) and the total N-terminal cleavage product (NE+NC) over the total proteins (NE+NC+P), respectively. 0-hour splicing and cleavage product values were subtracted from respective sample values for baseline corrections. The results were analysed using one-way ANOVA and plotted using GraphPad Prism version 5.01 for Windows, GraphPad Software, San Diego, CA, U.S.A., www.graphpad.com.

### 2.8 Western blot analysis ^8^

For Western blot analysis, an anti-His antibody was utilized to confirm the presence of splicing and N-terminal cleavage products. The test proteins were transferred from the 5-10% SDS-PAGE to the PVDF membrane (Millipore, IPVH 00010) at a voltage of 10V, overnight at 4°C. Subsequently, the blot was blocked with 5% skim milk and incubated overnight at 4°C. Then, the membrane was washed once with 1× TBST (Tris-buffered saline with Tween-20) followed by incubation with a primary anti-His antibody (abgenex-32-6116) at 1:5000 dilution overnight at 4°C. In the next step, the membrane was washed thrice for 5 minutes, with 1×TBST and incubated with a secondary antibody (anti-mouse, abgenex 11-301) at a dilution of 1:6000, for 6 hours at 25°C. To enhance detection of the N-extein band, a higher dilution of the primary antibody (1:2500) and secondary antibody (1:4000) were used. Finally, the blots were washed with 1× TBST and developed using Enhanced Chemiluminescence Substrate (ECL) (abcam, ab65628) to visualize the protein bands and images were captured using in-house ImageQuant^TM^ LAS 500 facility (GE Healthcare).

### 2.9 Anti-mycobacterial study

#### 2.9.1 Preparation of Mtb cells ^65^

Different mycobacterial cells [Drug sensitive H37Rv (ATCC-27294), H37Ra (ATCC-25177), *M.sm* (ATCC-19420) and clinically isolated MDR *Mtb* isolates obtained from National Reference Laboratory (NRL), ICMR-Regional Medical Research Centre (RMRC-ICMR), Bhubaneswar, India] were grown in Middlebrook 7H9 broth (Merck-M0178) supplemented with Middlebrook oleic acid, albumin, dextrose, and catalase (OADC) (Himedia-FD018). CFU calculations were made by matching the turbidity of inoculum with McFarland standard (Himedia-R092) against a black background. 10^6^ CFU of mycobacterial cells were used for subsequent experiments.

#### 2.9.2 Alamar blue assay ^66–68^

10^6^ CFU of mycobacterial cells were incubated with different concentrations of ZnONPs (0.5 ug/ml to 50 ug/ml) in 96-well microtiter plates. For a comparative study, cells were also incubated with anti-TB drugs Isoniazid (INH), Rifampicin (RIF), Moxifloxacin (MXF), and Levofloxacin (LFX) to evaluate their activity towards H37Rv strain and multidrug-resistant (MDR) *Mtb* isolates. The plates were sealed and incubated at 37°C for 14 days. Later, 10% (v/v) solution of Alamar Blue reagent (SRL-42650) was added to each well and the plates were incubated at 37°C for 24 hours. The anti-mycobacterial efficacy of each compound was evaluated based on the observed colour change. Live cells were identified by transition of colour from blue to pink whereas dead cells remained blue. The minimum inhibitory concentration (MIC) for ZnONPs and anti-TB drugs were determined as the lowest concentration at which cell viability was inhibited partially, resulting in a purple colour.

#### 2.9.3 Spread plate method to determine Minimum Bactericidal Concentration (MBC) ^69^

Mycobacterial cells at 10^6^ CFU were incubated with different concentrations of ZnONPs (0.5 μg/ml to 50 μg/ml) for 14 days in 96 well microtiter plates. Then, sample (1:1000 dilution) from each well was evenly spread onto respective 7H10 agar (Himedia-M199) plate using a sterile spreader. The plates were incubated at 37°C for 21days to grow mycobacterial colonies. Subsequently, colonies were counted by colony counting method and MBC was determined as the lowest concentration of the test compounds at which no colony growth was visualised on the respective agar plates. Controls plates included mycobacterial cells without NPs treatment.

#### 2.9.4 Scanning Electron Microscopic (SEM) analysis ^70^

10^6^ CFU of mycobacterial cells were incubated with different concentrations (0.5 μg/ml,17 ug/ml, 26 μg/ml, and 50 μg/ml) of ZnONPs in 7H9 broth medium for 7 days. Then, cells were harvested, fixed with 2.5% glutaraldehyde, and dehydrated with a series of ethanol treatment (30%, 60%, 70%, 80%, and 90%). Finally, cells were treated with 100% ethanol for 30 min. Then gold-palladium particles were sputtered over the mycobacterial cells by an automated sputtering machine and visualized via SEM (Hitachi, S-3400N), CIF, OUAT, Bhubaneswar.

#### 2.9.5 Animal experiments ^66^

Female BALB/c mice, aged 6 to 8 weeks and weighing each 25 ± 2 g, were used for the study. Mice were randomly divided into three groups, each consisting of 6 to 8 mice. The animals were injected intravenously with 5 x 10^6^ colony-forming units (CFU) of *Mycobacterium tuberculosis* (strain H37Ra) over a period of 4 days. Subsequently, ZnONPs were orally administered to the infected mice at different concentrations (26 and 50 µg/ml) for 7 days post-infection. Rifampicin alone, given orally at a dose of 15 mg/kg to the infected mice was used as a positive control, while mice who were given oral 1×PBS served as the negative control. After 18 days of infection, the mice were euthanized under 5% inhaled isoflurane anaesthesia (Sigma-792632), followed by cervical dislocation. Then, their lungs and spleen were isolated and processed for histopathological examination and estimation of mycobacterial load via CFU analysis. A portion of the lung was fixed in 4% paraformaldehyde for 48 hours and then embedded in paraffin to prepare sections for haematoxylin and eosin staining. Next, lung and spleen homogenates in 1×PBS were plated onto Middlebrook 7H11 agar, supplemented with OADC (Himedia-FD018) and BBL MGIT PANTA antibiotics (Catalogue #245114, BD, United States), to quantify the bacterial burden. Culture plates were incubated for 3 to 4 weeks at 37°C, followed by calculation of CFU by colony counting methods.

### 2.10 Statistical analysis

For each method, numerical data from three to six independent sets of experiments were analysed using GraphPad Prism 6.0 (La Jolla, CA, United States) and are presented as mean ± 1 standard deviation or standard error of the mean. Statistical significance between different groups was assessed using one-way ANOVA, with significance denoted by p < 0.05. Specific p-values of <0.05, <0.01, <0.001, and <0.0001 were indicated as *, **, ***, and ****, respectively.

## 3. Results and discussions

### 3.1 ZnONPs interact with Mtb SufB precursor protein

Previous studies have reported interaction of nanoparticles with biological molecules such as proteins and DNA ^71^. Before examining the role of ZnONPs on the splicing activity, we evaluated possible interaction between ZnONPs and *Mtb* SufB precursor protein. Through a combination of analytical techniques such as UV-visible spectroscopy, Bradford analysis, and transmission electron microscopy (TEM), we have evaluated ZnONPs∼protein binding activity (Figure 1).

**Figure 1.**
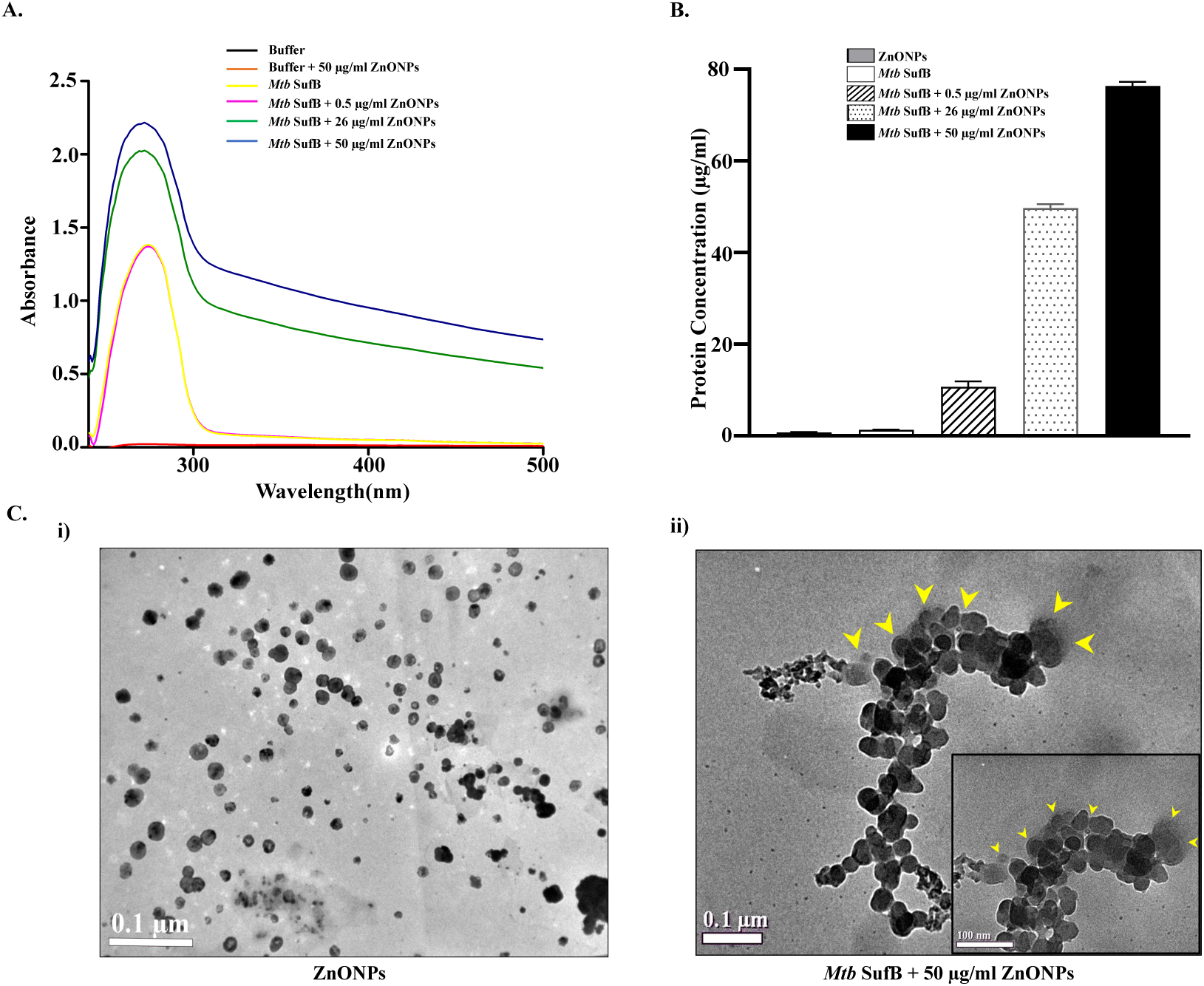
Evaluation of ZnONPs interaction with *Mtb* SufB. **A)** UV–Visible spectroscopy: *Mtb* SufB precursor protein was treated with ZnONPs (0.5 µg/ml, 26 µg/ml, 50 µg/ml), and sample absorbance was measured by scanning over 200–500 nm wavelength. **B)** Bradford Assay: Denatured test proteins were renatured *in vitro* in the presence of varied ZnONPs concentration for a period of 4 hrs at 25^0^C. Then, test samples were processed, and the concentration of adsorbed protein was determined using Bradford reagent. **C)** TEM Analysis: Images showing. (i) discrete ZnONPs and (ii) protein corona formation on the surface of ZnONPs aggregates after interaction with *Mtb* SufB (yellow arrows). The inset focuses on the protein shell around the ZnONPs.

Initially, cytotoxicity of the ZnONPs was assessed via MTT assay using human embryonic kidney cell line, HEK239T cells, since potentially harmful NPs are mostly excreted by kidney through urine ^72–74^. It was demonstrated that up to a concentration of 50 µg/ml, ZnONPs exhibited no cytotoxicity against HEK239T cells (Figure S1, Supplementary Information). This is further supported by a prior research which has shown similar results ^75^. Consequently, we decided to explore SufB interaction with various ZnONPs concentrations up to this limit (0.5, 26, and 50 µg/ml). UV–Visible spectra for both the *Mtb* SufB precursor protein and ZnONPs-SufB complexes were recorded by scanning over the visible range (200 to 800 nm), as depicted in Figure 1A. Buffer alone and buffer along with nanoparticles (50 µg/ml) were used for baseline correction and as the reference blank respectively. An observable broadening and blue shift were differentiated in the absorbance spectra of the *Mtb* SufB protein following interaction with increasing concentrations of ZnONPs, when compared to the spectra of protein solubilised in buffer alone. This observation suggests the binding of ZnONPs with the *Mtb* SufB precursor protein over test concentration range.

To further validate such interaction, we utilized Bradford assay, where *Mtb* SufB precursor protein was refolded along with varying concentrations of ZnONPs (0.5, 26, and 50 µg/ml), causing adsorption of proteins onto the NPs surface. After isolating ZnONPs-protein complexes, quantification of bound proteins was done using Bradford assay. As illustrated in Figure 1B, an increase in nanoparticle concentration correlated with a proportional increase in the bound proteins. This result further supported the interaction between ZnONPs and *Mtb* SufB protein.

The formation of a protein corona on NPs surfaces was investigated through TEM analysis, as shown in Fig 1C. While standalone ZnONPs exhibited discrete distribution [1C(i)], NPs (50 µg/ml)-treated protein sample displayed a distinct feature. A thin, transparent layer encasing the NPs was evident in the treated sample (yellow arrows) as seen in the Figures 1C(ii and iii). This is indicative of the protein corona formed on the surface of the ZnONPs upon interaction with the *Mtb* SufB protein ^76^.

DLS measurements demonstrated a significant increase in hydrodynamic diameter following interaction of ZnONPs (26 µg/ml and 50 µg/ml) with *Mtb* SufB compared to controls (Buffer; Buffer with *Mtb* SufB; Buffer with ZnONPs) (Figure S2A, Table S1, Supplementary Information)^77^. Concomitant zeta potential analysis revealed a charge inversion from +5.1 mV (bare ZnONPs) to -4 mV for the ZnONPs-SufB complex (Figure S2B, Table S1, Supplementary Information)^77,78^. This transition from net positive to net negative surface charge suggests a protein-nanoparticle binding event. Therefore, the observed changes in both hydrodynamic size and surface charge distribution shows evidence for ZnONPs-SufB interaction.

The comprehensive assessment involving UV-visible spectroscopy, Bradford analysis, DLS, zeta potential analysis and TEM, successfully indicated the binding activity and protein corona formation following ZnONPs interaction with *Mtb* SufB.

### 3.2 ZnONPs inhibit generation of functional SufB protein

The SUF system provides an unique pathway for [Fe-S] cluster biosynthesis and *Mtb* survival during intracellular stress like iron deprivation ^79^. In such environments, the functionality of the SUF complex relies exclusively on the splicing of *Mtb* SufB precursor to generate active SufB protein ^13^. Hence, splicing regulation under such physiological stress can lead to potential anti-TB drug development strategies by influencing mycobacterial cell viability. Although metals are known inhibitors of intein splicing ^42,45,46^, NPs mediated splicing inhibition has not been reported yet. Further, higher reactivity of nanoparticles in a biological system has prompted us to evaluate ZnONPs and *Mtb* SufB interactions with possible effects on SufB precursor splicing.

*In vitro* splicing assay was conducted both in the presence and absence of ZnONPs, as detailed in the methods section. Since, MTT assay demonstrated that HEK239T cells (human embryonic kidney cell lines) were tolerant to ZnONPs up to a concentration of 50 µg/ml (Figure S1, Supplementary Information), splicing efficiency was evaluated using NPs concentrations up to this limit. Figure 2A illustrates different structural domains of *Mtb* SufB precursor protein; N-extein (29.9 kDa), Intein (40.2 kDa), and C-extein (25.8 kDa). The details of *Mtb* SufB splicing and N-terminal cleavage reactions are published elsewhere ^7^. SufB precursor undergoes splicing reactions to generate ligated exteins (LE), and intein (I), whereas N-terminal cleavage reactions yield N-terminal cleavage product (NC, 65.2 kDa), and N-extein (NE) (Figure 2A). Figures 2B(i) and 2B(ii) depict the concentration-dependent inhibition of these reactions by ZnONPs, providing crucial insights into their biological role.

**Figure 2.**
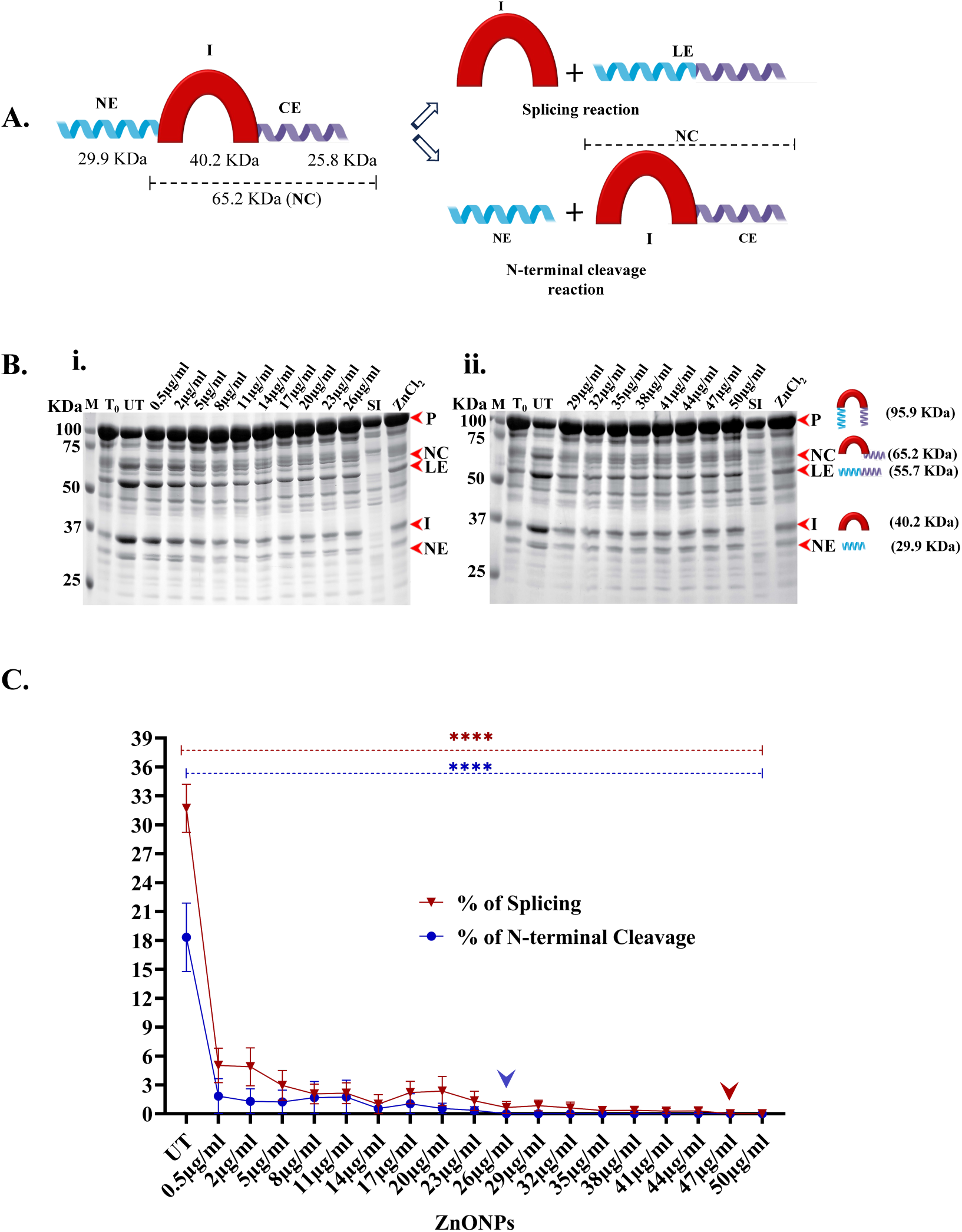
Effect of ZnONPs on Mtb SufB precursor splicing and N-terminal cleavage reactions. **(A)** Schematic diagram representing different structural domains of *Mtb* SufB precursor protein and splicing & N-terminal cleavage reactions of the precursor protein generating different protein products. **(B)** Products from *in vitro* protein refolding experiment were resolved through 5–10% gradient SDS-PAGE. (T0): splicing and cleavage reactions at 0 h, (UT): untreated protein sample; and Lanes 4-13 show protein products induced by varied concentrations ZnONPs: (i) (0.5 µg/ml -26 µg/ml) (ii) (29 µg/ml to 50 µg/ml), Lane 14 (SI): Splicing inactive SufB double mutant (Cys1Ala/Asn359Ala) is used as a negative control for splicing, Lane 15: ZnCl2 (2 mM) was used as a positive control, since it is a known inhibitor of *Mtb* SufB splicing reactions ^8^. **(C)** Line plots showing the quantitative results of splicing and N-terminal cleavage reaction inhibition. Values were extracted from Figure 2(B) and plotted after densitometric analyses using GelQuant.Net biochemical solutions. 0h splicing, and cleavage values were subtracted from each sample as baseline correction. The red arrow and blue arrow denote complete inhibition of SufB splicing and N-terminal cleavage reactions by ZnONPs at concentrations of 47 µg/ml and 26 µg/ml, respectively. Error bars represent (±1) SEM from (n=3) three independent sets of experiments (p<0.0001). P: precursor, NC: N-terminal cleavage product, LE: ligated extein, I: intein, and NE: N-extein.

The samples were analyzed through gradient SDS-PAGE, and the bands were quantified using the GelQuant.NET software. Our findings indicated a substantial inhibition of *Mtb* SufB precursor splicing (p<0.0001) over all the test concentration range, with discernible effects commencing at a ZnONPs concentration of 0.5 µg/ml. Remarkably, a complete inhibition of splicing was observed at a concentration of 47 µg/ml [Figures 2B(ii) and C(i)]. These findings suggest that ZnONPs exert a dose-dependent inhibition of splicing in *Mtb* SufB precursor protein which could lead to critical consequences such as loss of functionality of this protein in a cellular environment.

Similarly, the N-terminal cleavage reaction of *Mtb* SufB precursor was notably inhibited (p<0.0001) in the presence of ZnONPs. A reduction in the cleavage activity was observed at the minimal tested concentration of 0.5 µg/ml, culminating in a complete inhibition at a concentration of 26 µg/ml [Figures 2B(i) and C(ii)]. Thus, ZnONPs exert a potent inhibitory effect on the *Mtb* SufB splicing and N-terminal cleavage reactions, underscoring their potential interference with SufB-associated cellular processes. These findings also highlight the significant biological implications of ZnONPs on *Mtb* survival within host cells. To substantiate these findings, we performed western blot analysis to validate identity of protein products (Figure S3, Supplementary Information), which further confirmed the inhibitory role of ZnONPs on the splicing and N-terminal cleavage reactions of SufB. Additional intermediate concentrations of ZnONPs were used to check the robustness of such effects on the splicing and cleavage activity of the precursor protein (Figure S4, Supplementary Information).

Current research presents evidence that ZnONPs interact with the *Mtb* SufB precursor protein, leading to a concentration-dependent inhibition of both splicing and N-terminal cleavage reactions. These findings also emphasize on the notable impact ZnONPs could have on the essential cellular processes linked to *Mtb* SufB protein, and their potential implications in biological systems. Specifically, it calls for further exploration into the influence of ZnONPs on SufB splicing, correlating this process to the growth and viability of *Mtb*.

### 3.3 ZnONPs inhibit the growth and viability of Mycobacterial cells

Since ZnONPs were shown to interact with the SufB precursor protein, and caused inhibition of protein splicing, it was imperative that this may retard the growth and viability of *Mtb* by blocking the generation of an essential protein SufB. Both H37Rv cells and MDR *Mtb* isolates, were cultured in a nutrient medium along with various concentrations (0.5 µg/ml∼50 µg/ml) of ZnONPs, as discussed in the methods section. The test concentration range for the ZnONPs were chosen based on their activity range as outlined in Figures 2B and 2C. Further, up to a concentration of 50 µg/ml, ZnONPs exhibited no cytotoxicity against HEK239T cells (Figure S1, Supplementary Information). Besides, ZnONPs at a concentration of ≥50 µg/ml was also found to be toxic for human cells, elucidated by earlier research ^80^.

We performed Alamar blue assay, a well-established metric for cell viability, which yielded data on the inhibitory roles of ZnONPs on the growth and survival of H37Rv and MDR *Mtb* cells. The Alamar blue dye, initially blue, undergoes reduction in viable cells, resulting in a colour change to pink, whereas persistence of the blue colour indicates lack of viable cells ^81^. ZnONPs at 0.5 μg/ml and 2 μg/ml did not have any effect H37Rv cells. 5 μg/ml was identified as the minimum inhibitory concentration (MIC) for ZnONPs at which H37Rv cells turned purple due to partial loss of cell viability (Figure 3A). A complete loss of cell viability, and retention of blue colour was noted at a concentration of 17 μg/ml of ZnONPs.

**Figure 3.**
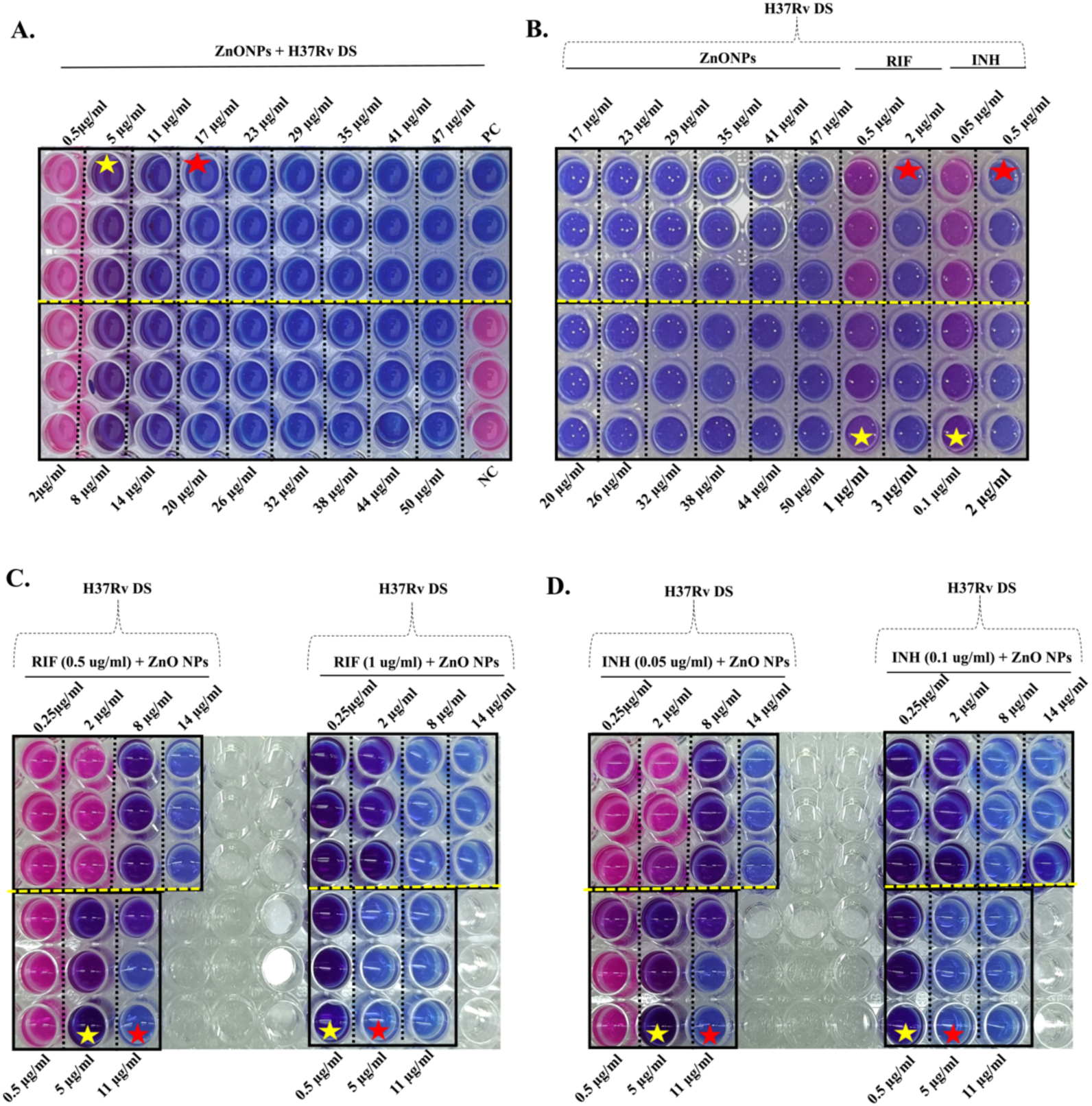
Alamar blue assay to determine the minimum inhibitory concentrations (MIC) of ZnONPs towards H37Rv *Mtb* cells. (A) Mycobacterial cells were incubated in a 96 well plate, in presence of varied concentrations (0.5 to 50 µg/ml) of NPs for 14 days, followed by addition of Alamar Blue reagent. Further incubation at 37°C for 24 hours, led to colour change in the wells. MIC of ZnONPs was identified as 5 μg/ml where transition of blue to purple colour occurred suggesting partial loss of cell viability among the cells. Persistence of blue colour correlating to complete loss of viable cell population was seen at 17 μg/ml of ZnONPs. Pink cells refer to healthy viable cells resisting inhibitory effect of ZnONPs. (B) Comparison of the activity of first line anti-TB drugs; INH & RIF and varied concentration of ZnONPs. All the cells were rendered non-viable (blue) at a concentration of 17 (g/ml for the NPs, 2 μg/ml for RIF, 0.5 μg/ml for INH, and at respective higher concentrations. The MIC for RIF & INH as shown in this study coincided with the known values of 1 and 0.1 μg/ml respectively ^82^. Synergistic activity of ZnONPs and anti-TB drugs; (C) RIF & (D) INH. 10^6^ CFU of bacterial cells were grown in presence of varied concentrations of ZnONPs and Drugs. A combination of RIF (0.5 μg/ml)+ ZnONPs (11 (g/ml) & RIF (1 μg/ml)+ ZnONPs (5 μg/ml) successfully yielded non-viable cells (blue). Likewise, INH (0.05(g/ml) + ZnONPs (11 μg/ml) & INH (0.1 μg/ml) + ZnONPs (5 μg/ml) combinations resulted in non-proliferating cells (blue). Red stars-complete loss of viability & Yellow stars-partial loss of viable cells (purple) at a lower concentration of NPs. DS: Drug sensitive H37Rv *Mtb*, RIF: Rifampicin, INH: Isoniazid.

Next, a comparative study was conducted to assess the efficacy of ZnONPs in tandem with first-line anti-TB drugs such as Rifampicin (RIF) and Isoniazid (INH) for which the Minimum Inhibitory Concentrations (MICs) and Minimum bactericidal Concentrations (MBC) are well-documented ^82,83^. RIF and INH, at known MICs of 1 μg/ml and 0.1 μg/ml respectively, resulted in partial loss of mycobacterial cell viability (purple colour) in the respective wells. There was a complete loss of viable cell population (blue colour) at 2 μg/ml of RIF, and 0.5 μg/ml of INH, comparable to the activity of ZnONPs at 17 μg/ml (Figure 3B). All the cells were rendered non-viable at concentrations higher than these values.

Figures 3C and 3D depict the synergistic activity of ZnONPs and first line anti-TB drugs; RIF & INH. A combination of RIF (0.5 μg/ml) + ZnONPs (11 μg/ml) & RIF (1 μg/ml) + ZnONPs (5 μg/ml) successfully yielded non-viable cells (blue). Likewise, INH (0.05 μg/ml) + ZnONPs (11 μg/ml) & INH (0.1 μg/ml) + ZnONPs (5 μg/ml) combinations resulted in non-proliferating cells (blue). Thus, both RIF and INH at their respective MIC and half reduced MIC values could completely abolish mycobacterial cell viability when added together with ZnONPs. These results emphasize on the diminution in effective concentration for the conventional anti-TB drugs (INH & RIF) in a combination with ZnONPs.

MIC values of ZnONPs towards multidrug-resistant (MDR) *Mtb* isolates were found to be higher at 14 μg/ml (Figure 4A). The MBC values for the ZnONPs [17 µg/ml for drug-sensitive H37Rv and 23 µg/ml for multidrug-resistant (MDR) *Mtb* isolates] were further confirmed through agar spread plate method as shown in Supplementary Information (Figure S5, Supplementary Information). Although MDR cells were rendered nonviable at 20 μg/ml of ZnONPs (Figure 4A), based on the data obtained from agar spread plate method (Figure S5, Supplementary Information), for the remaining experiments 23 µg/ml was considered as the MBC for ZnONPs towards MDR *Mtb* cells.

**Figure 4.**
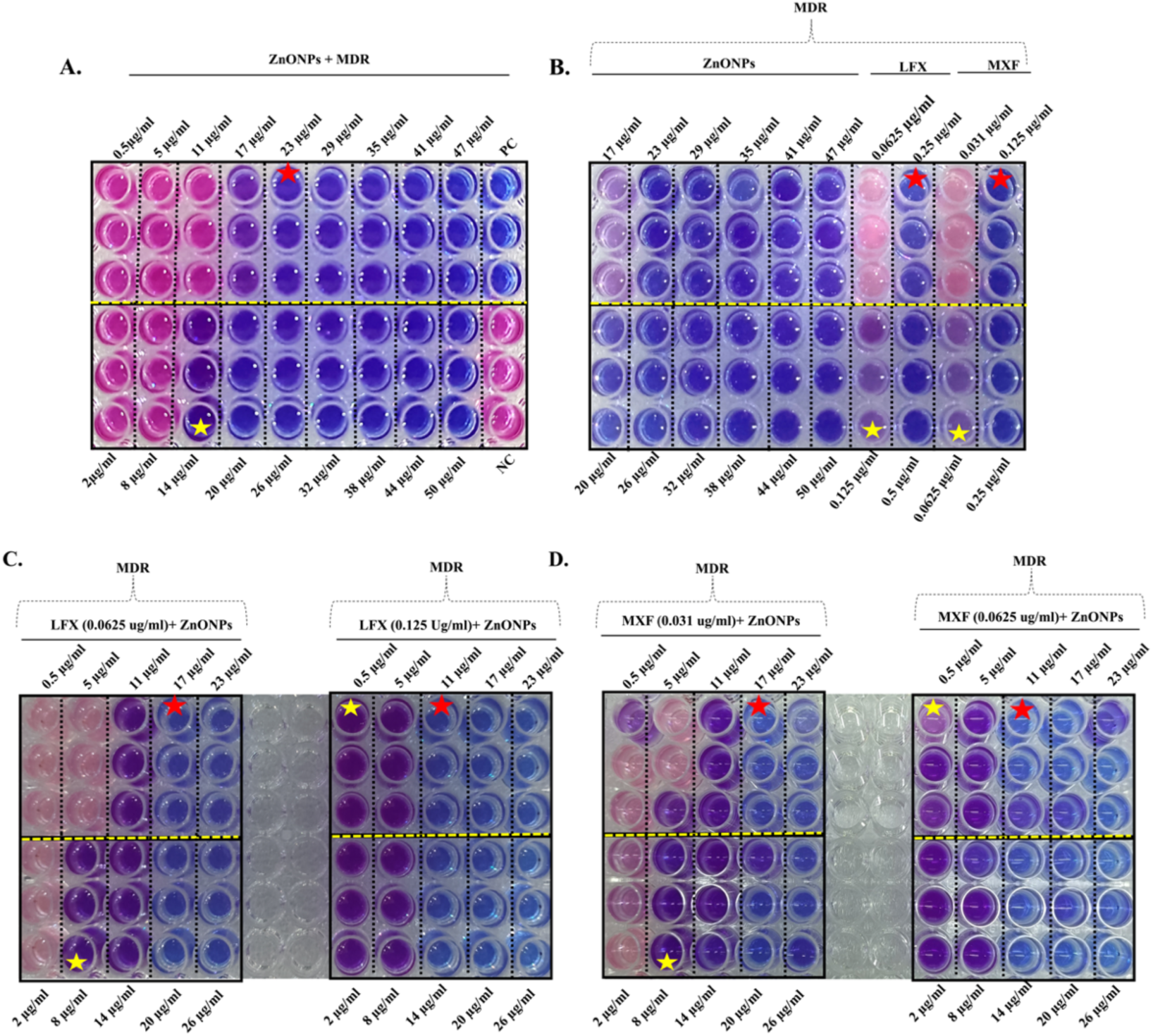
Alamar blue assay to determine the MIC of ZnONPs towards MDR *Mtb* cells. (A) Mycobacterial cells were incubated in a 96 well plate, in presence of varied concentrations (0.5 to 50 µg/ml) ZnONPs for 14 days, followed by addition of Alamar Blue reagent. Further incubation at 37°C for 24 hours, led to colour change in the wells. MIC of ZnONPs was identified as 14 (g/ml where transition of blue to purple colour occurred suggesting partial loss of cell viability among the cells. In contrast, persistence of a blue colour due to complete loss of viable cell population was seen at 23 (g/ml of ZnONPs. Pink cells refer to healthy viable cells resisting inhibitory effect of ZnONPs. (B) Comparison of the activity of anti-TB drugs; Levofloxacin (LFX) & Moxifloxacin (MXF) and varied concentration of ZnONPs. All the cells were rendered non-viable (blue) at a concentration of 23 (g/ml for the ZnONPs and in presence of LFX (0.25 (g/ml), MXF (0.125 (g/ml) and at higher concentration. The MIC for LFX & MXF as shown in this study coincided with the known values of 0.125 and 0.0625 (g/ml respectively ^84,85^. Synergistic activity of ZnONPs and anti-TB drugs; (C) LFX & (D) MXF. 10^6^ CFU of bacterial cells were grown in presence of varied concentrations of ZnONPs and the drugs. A combination of LFX (0.0625 (g/ml) + ZnONPs (17 (g/ml) & LFX (0.125 (g/ml) + ZnONPs (11 (g/ml) successfully yielded non-viable cells (blue). Likewise, MXF (0.031 (g/ml) + ZnONPs (17 (g/ml) & MXF (0.0625 (g/ml) + ZnONPs (11 (g/ml) combinations resulted in non-proliferating cells (blue). Red stars-complete loss of viability & Yellow stars-partial loss of viable cells (purple) at a lower concentration of NPs. MDR: Multidrug-resistant, LFX: Levofloxacin, MXF: Moxifloxacin.

In the case of MDR isolates, the anti-TB drugs Moxifloxacin (MXF) and Levofloxacin (LFX), with known MICs of 0.0625 μg/ml and 0.125 μg/ml respectively^84,85^, were compared with the activity of the ZnONPs (Figure 4B). There was a complete loss of viable cell population (blue colour) at LFX (0.25 μg/ml) / MXF (0.125 μg/ml) and their activity was comparable to the effect of ZnONPs at 23 μg/ml (MBC) (Figure 4B). All the cells were rendered non-viable (blue) at concentrations higher than these values for the test compounds. Thus, the antimycobacterial activity of ZnONPs towards MDR *Mtb* isolates was comparable to that of anti-TB drugs MXF and LFX at their respective MIC and higher values. This underscores the potential of ZnONPs as a potent alternative for managing *Mtb* infections

Next, the synergistic activity of ZnONPs along with LFX and MXF was evaluated towards MDR *Mtb* isolates (Figures 4C and 4D). A combination of LFX (0.0625 μg/ml)+ ZnONPs (17 μg/ml) and LFX (0.125 μg/ml)+ ZnONPs (11 μg/ml) resulted in non-viable cells (blue). A similar observation was made for, MXF (0.031 μg/ml) + ZnONPs (17 μg/ml) and MXF (0.0625 μg/ml)+ ZnONPs (11 μg/ml) combinations towards the viability of MDR cells.

Thus, both LXF and MXF at their respective MIC and half reduced MIC values could completely abolish mycobacterial cell viability when added together with ZnONPs. In toto, these results suggest that ZnONPs exhibit anti-mycobacterial properties against both H37Rv and MDR *Mtb* cells, indicating their potential utility as a supportive therapy in the fight against TB.

On the whole, from the results of *in vitro* splicing and Alamar blue assays, ZnONPs-mediated inhibition of mycobacterial growth could be attributed to splicing attenuation induced by interactions between ZnONPs and *Mtb* SufB precursor protein. By targeting essential proteins like the splicing of SufB, ZnONPs are likely to disrupt vital cellular processes in the mycobacteria, leading to growth inhibition and loss of cell viability. To further correlate these events, Alamar blue assay was performed to examine the effect of ZnONPs (0.5 ug/ml to 50 ug/ml) on the growth and viability of *Mycobacterium smegmatis* (*M. sm*), in which SufB polypeptide chain lacks an intein sequence (Figure S6, Supplementary Information) ^86^. Live viable cells (pink) (Figure S6, Supplementary Information), indicated no effect of ZnONPs (0.5 µg/ml to 50 µg/ml) on *M. sm* viability, possibly due to lack of regulatory influence on SufB splicing and cleavage reactions. This provided further evidence to support loss of *Mtb* cell viability (alamar blue assay) as a result of ZnONPs mediated inhibition of *Mtb* SufB splicing and cleavage reactions, although additional mechanisms such as NPs-mediated membrane damage cannot be ruled out.

Furthermore, our investigation into combinatorial therapy, integrating lower effective concentrations of known anti-TB drugs alongside ZnONPs, revealed a notable inhibitory effect on the growth and viability of *Mycobacterium tuberculosis* (*Mtb*) cells. Particularly noteworthy was the complete loss of cell viability observed at concentrations half of the MIC values for the individual drugs along with a reduced effective concentration for the NPs (Figures 3C, 3D, 4C, and 4D). As shown in Figure 3C, when the concentration for RIF was reduced to 0.5 µg/ml (half of MIC), along with 11 µg/ml of ZnONPs, this led to no observable viable growth of H37Rv *Mtb*. Similarly, INH at a concentration of 0.05 ug/ml (half of MIC) completely blocked viability of mycobacterial cells in presence of lower dose of ZnONPs (11 µg/ml) (Figure 3D). In another approach, when RIF and INH were added at their respective MIC (1 µg/ml and 0.1 µg/ml respectively), in presence of MIC of ZnONPs (5 µg/ml), no viable cells were observed (Figures 3C, 3D). Similarly, in the case of MDR cells, when LFX and MXF were added at their MIC (0.125 and 0.0625 µg/ml respectively)^84,85^, a reduction for ZnONPs MIC occurred from 14 ug/ml to 0.5 ug/ml, with no growth observed at 11 µg/ml (Figures 4C, 4D).

This strategy demonstrates the potential to achieve significant inhibition of H37Rv as well MDR *Mtb* cells at markedly lower concentrations of conventional anti-TB drugs, with the potential to mitigate the cytotoxic effects associated with prolonged drug administration. The observed synergy between lower doses of existing drugs and ZnONPs presents a promising avenue for enhancing the efficacy of TB management while minimizing potential adverse effects.

### 3.4 ZnONPs exhibit bactericidal effects on drug-sensitive and MDR Mtb cells

Current work validated the interaction between ZnONPs and *Mtb* SufB, which influenced cell viability, and this can be attributed to the production of a non-functional SufB protein via splicing inhibition, a vital component for *Mtb* survival ^11,13^. The observed decrease in viability could also be associated with a cumulative effect of mycobacterial membrane damage activity induced by ZnONPs ^40^. To substantiate this hypothesis, we performed a scanning electron microscopy (SEM) analysis, employing various ZnONPs concentrations; a lower concentration of 0.5 μg/ml, two effective concentrations of 17 and 23 μg/ml (MBC for ZnONPs towards H37Rv and MDR *Mtb* cells), and 50 μg/ml, the upper limit for toxicity. These concentrations were selected to cover the effective range of ZnONPs for *in vitro* splicing studies, and MBC values identified by Alamar blue assay & spread plate method.

Treatment with ZnONPs at a concentration of 0.5 μg/ml induced minor alterations in the bacterial morphology. However, exposure to concentrations of 17 μg/ml, 23 μg/ml, and 50 μg/ml led to extensive membrane disruption, cell-fusion, and cytoplasmic leakage accompanied by a complete loss of cellular morphology (Figures 5A and 5B). Thus, SEM analysis offers substantial evidence of the membrane-damaging activity of ZnONPs towards H37Rv and MDR *Mtb* isolates. This uniform activity of ZnONPs against both drug-sensitive and MDR *Mtb* cells validates our previous observations on antimycobacterial effectiveness detected via Alamar blue assay. This result is particularly significant in the context of MDR-TB, where conventional antibiotic treatments encounter substantial obstacles.

**Figure 5.**
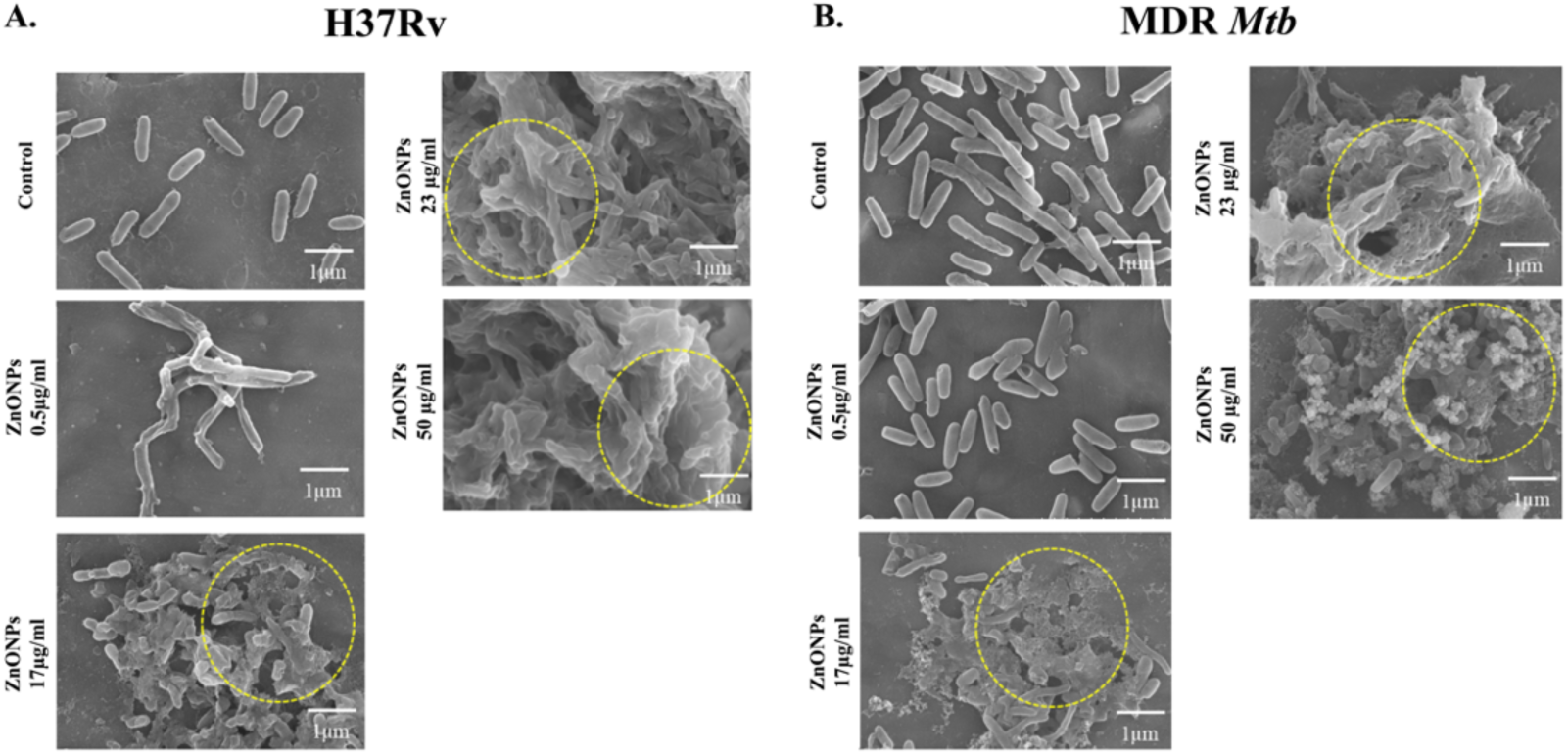
SEM analysis displaying bactericidal role of ZnONPs on *Mycobacterium tuberculosis (Mtb)* cells. **(A)** H37Rv **(B)** MDR *Mtb* isolates. Control (untreated) mycobacterial cells were uniform looking, without any changes in the cell morphology or any evidence of membrane damage. Mild alterations in cellular morphology were noted for both H37Rv, and MDR *Mtb* cells in presence of 0.5 µg/ml of ZnONPs. Dose dependent increase in the severity of cell membrane damage, cell fusion, loss of cellular architecture, were identified among the **(A)** H37Rv and **(B)** MDR *Mtb* isolates, across rest of the ZnONPs test concentration range (17 µg/ml, 23 µg/ml, and 50 µg/ml) (outlined by yellow circles). MDR: Multidrug-resistant *Mtb*

### 3.5 ZnONPs rescue Mtb induced changes in infected mice model

So far, *in vitro* data from this work suggested potent mycobactericidal activity of ZnONPs against H37Rv and MDR *Mtb* isolates, possibly by inhibiting the splicing of essential protein *Mtb* SufB, leading to loss of viability and membrane damage. To further assess this activity in *in vivo* setting, we conducted experiments in mice model.

The utilization of *Mtb* H37Ra (H37Rv attenuated strain)-infected mice serves as an established *in vivo* model to evaluate TB infection ^87^. Both H37Ra and H37Rv *Mtb* strains are inhibited by similar concentrations of anti-TB drugs as suggested by an earlier work ^88^. Hence, the efficacy of ZnONPs-drug combination and recovery from the infection was further tested in H37Ra-infected mice.

BALB/6 mice were infected with *Mtb* H37Ra; 1×PBS (placebo), ZnONPs, and drug (RIF) administration commenced four days post-infection and continued for a total of seven days, with mice sacrificed on day 18 post-infection for subsequent histopathological and bacterial burden analysis (experimental design elucidated in Figure 6A). Administration of RIF dosages is based on earlier works published elsewhere ^89–91^. Our findings revealed a strong anti-*Mtb* effect of ZnONPs, as evidenced by the analysis of the gross morphology of spleen, histopathological analysis and bacterial burden assay.

**Figure 6.**
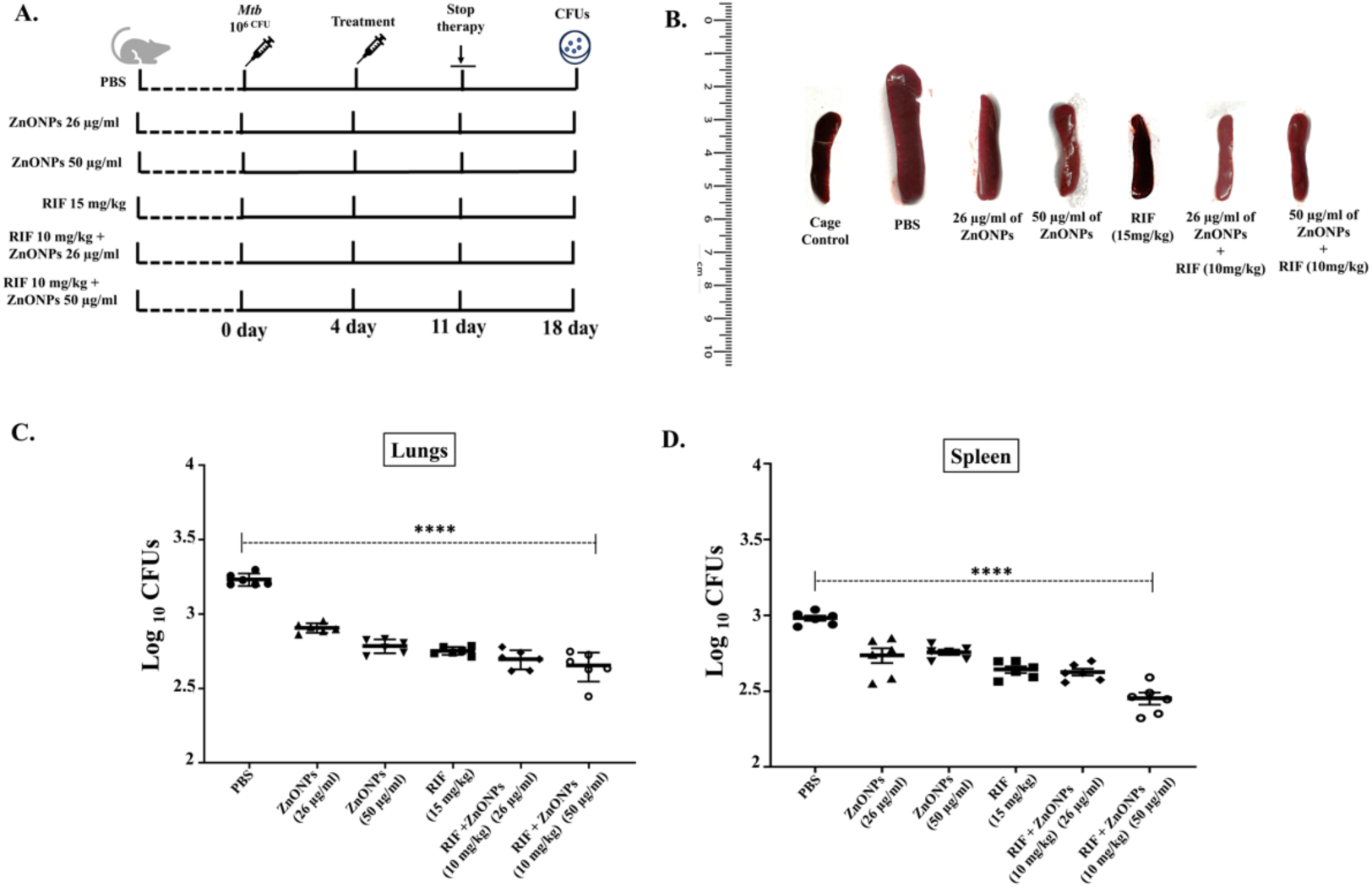
*In vivo* antimycobacterial effects of ZnONPs in H37Ra infected mice model. **(A)** Schematic representation of the experimental design in BALB/c mice (n = 6 in each group). Mice were infected with H37Ra; 1×PBS (placebo), NPs and drug administration commenced four days post-infection and continued for a total of seven days, with mice sacrificed on day 18^th^ post-infection. **(B)** Significant splenomegaly was noticed in the infected mouse which had received 1×PBS only. This was not observed in the infected mice, who were given RIF (15 mg/kg) alone, and combinations of RIF (10 mg/kg) + ZnONPs (26 µg/ml), and RIF (10 mg/kg) + ZnONPs (50 µg/ml), where the spleen size was comparable to the normal sized spleen in healthy cage control mouse. The splenomegaly was partially rescued in mice who had received ZnONPs (26 or 50 µg/ml) alone. Bacterial burden in the **(C)** lungs, and **(D)** spleen of the infected mice was determined by plating the organ lysate on the 7H10 plates after treatment with ZnONPs or 1*×*PBS for 7 days. Significant reduction in mycobacterial loads was observed in mice groups treated with a combination of RIF (10 mg/kg) + ZnONPs (26 µg/ml), and RIF (10 mg/kg) + ZnONPs (50 µg/ml), emphasizing the superior anti-TB activity of the ZnONPs-RIF combination as compared to RIF alone. Data are presented as mean ± 1 SD (p>0.0001) from values obtained from (n=6) mice.

A significant splenomegaly was observed in the infected mouse which had received 1×PBS as a placebo following infection with H37Ra strain ^92–96^. Resistance to this morphological effect was observed in infected mice, receiving RIF (15 mg/kg) alone, ZnONPs (26 and 50 µg/ml), RIF (10 mg/kg) + ZnONPs (26 µg/ml), and RIF (10 mg/kg) + ZnONPs (50 µg/ml). The splenomegaly following H37Ra infection in the aforementioned experimental groups was rescued significantly when ZnONPs was given as a solo treatment at concentration of 26 µg/ml, and 50 µg/ml. Further, the efficacy of RIF at 15 mg/kg was similar to a combination containing a reduced dosage of RIF at 10 mg/kg along with ZnONPs (26 µg/ml, and 50 µg/ml), where a complete rescue from splenomegaly was seen; comparable to the normal sized spleen in healthy cage control mouse (Figure 6B). Likewise, a substantial reduction in mycobacterial load in lungs and spleen were observed in similarly treated mice groups, emphasizing the superior anti-TB activity of the ZnONPs/RIF combination (Figures 6C and 6D). Bacterial loads were equivalently attenuated when RIF was administered solo at 15 mg/kg or in combination (RIF 10 mg/kg + ZnONPs 26 µg/ml or RIF 10 mg/kg + ZnONPs 50 µg/ml) at a lower concentration. This highlights the synergistic activity of these anti-*Mtb* compounds displaying a higher potential to manage mycobacterial infection.

Next, histopathological examination of infected lungs and spleen tissues provided additional evidence for the anti-mycobacterial efficacy of ZnONPs. Lungs sections from cage control healthy mice displayed clear lung fields without any evidence of inflammation [Figure 7A(a)]. Infected mice group administered with 1*×*PBS displayed distinct pathological changes in the lungs with noticeable inflammatory infiltrates, indicative of infection induced immune response [Figure 7A(b)] ^97^. When administered individually, ZnONPs (26 µg/ml), ZnONPs (50 µg/ml), and RIF (15 mg/kg) caused a partial reduction in inflammatory cells within the lungs [Figures 7A(c), (d), and (e)]. On the contrary, mice treated with a combination of ZnONPs (26 µg/ml) + RIF (10 mg/kg) and ZnONPs (50 µg/ml) + RIF (10 mg/kg) respectively, demonstrated no inflammatory infiltrations, suggesting their superior anti-mycobacterial action ([Figures 7A(f), and (g)].

**Figure 7.**
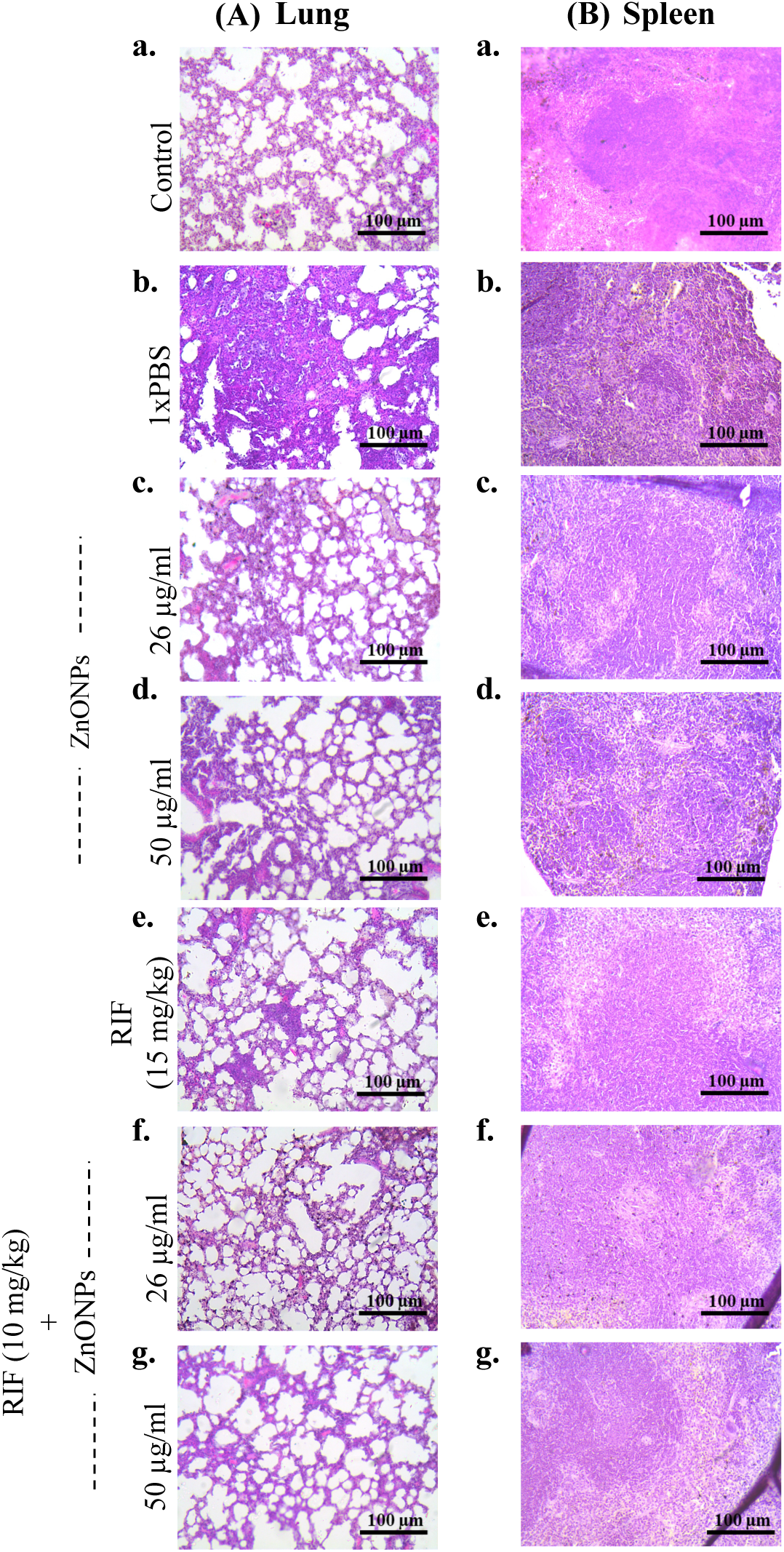
Histopathology of lungs and spleen from H37Ra infected mice (H37Rv attenuated strain), comparing the efficacy of ZnONPs, RIF, and ZnONPs-RIF combinations. **(A)** Lungs sections are stained by H&E staining and processed for visualization by microscopy. **(a)** Histology of uninfected lungs in healthy control mice. **(b-e)** represent tissue sections from infected mice who have received different solo treatments, (**b)** 1*×*PBS exhibiting prominent aggregates of inflammatory cells; (**c)** 26 ug/ml of ZnONPs, **(d)** 50 ug/ml of ZnONPs, and **(e)** RIF 15 mg/kg displaying partial reduction in immune cells infiltrates; **(f)** RIF (10 mg/kg) + ZnONPs (26 ug/ml), and **(g)** RIF (10 mg/kg) + ZnONPs (50 ug/ml) represent no inflammatory changes and restoration of normal lungs histology. **(B)** Tissue sections from spleen are stained by H&E staining and processed for visualization by microscopy. **(a)** Histology of spleen in uninfected healthy control mice showing well differentiated white and red pulp structures. **(b-e)** represent tissue sections from infected mice who have received different solo treatments; **(b)** 1*×*PBS exhibiting heavy immune cell aggregates obliterating white and red pulps; **(c)** 26 ug/ml of ZnONPs, **(d)** 50 ug/ml of ZnONPs, are displaying partial rescue from inflammatory infiltrates; **(e)** RIF 15 mg/kg, **(f)** RIF (10 mg/kg) + ZnONPs (26 ug/ml), and **(g)** RIF (10 mg/kg) + ZnONPs (50 ug/ml), either of these treatments when given to infected mice, restored normal spleen histology with no evidence of inflammation. Images represent different experimental groups of mice visualized under 10X magnification. H&E-Hematoxylin and eosin staining; RIF-Rifampicin

Similarly, inflammatory changes were noted within the white pulp of the infected splenic tissue, which is a critical component of the organ’s immune function ^98^. Healthy mice from the cage control group exhibited normal spleen histology with clear differentiation into white and red pulps [Figure 7B(a)] ^99^ whereas, infected mice treated with 1*×*PBS, depicted rich inflammatory infiltrates obscuring normal spleen histology [Figure 7B(b)] ^100^. A partial reduction in inflammation was observed, along with a restoration of the white and red pulp structures in the spleen of the mice treated with ZnONPs (26 µg/ml), and ZnONPs (50 µg/ml), respectively [Figures 7B(c), and (d)]. Notably, in the mice group that received RIF (15 mg/kg) alone or combination therapy [ZnONPs (26 µg/ml) + RIF (10 mg/kg) and ZnONPs (50 µg/ml) + RIF (10 mg/kg)], there were no inflammatory infiltrations, and distinct white and red pulp structures were evident ([Figures 7B(e), (f), and (g)]. Hence, the efficacy of the combinatorial approach with a reduced dose of RIF (10 mg/kg) was similar to RIF solo therapy at a higher concentration (15 mg/kg).

These observations support a potential therapeutic effect of the ZnONPs against *Mycobacterium tuberculosis* (*Mtb*). A concurrent mode of action combining, SufB splicing inhibition by ZnONPs along with inhibition of bacterial RNA polymerase due to RIF, and ZnONPs mediated membrane damage, possibly facilitates complete resolution of inflammation restoring normal spleen and lung histology ^101^. Taken together, these findings provide a strong foundation for further investigation into the mechanisms underlying the *in vivo* anti-mycobacterial activity of ZnONPs towards drug-sensitive as well as MDR TB. Additionally, long-term studies evaluating the safety and sustained efficacy of ZnONPs will be crucial for assessing their potential as a viable therapeutic option.

## 4. Conclusion

One of the major milestones of the United Nations Sustainable Development Goals (SDGs) is to end TB by 2030 ^102^. Besides aiming for accessible patient care with attenuated hospitalization rate, WHO also calls for “adoption to innovations” in TB management ^2^. The escalating challenge of drug-resistant TB, necessitates the exploration of novel therapeutic strategies such as nanomedicine. Current work investigates into the mycobactericidal role of the ZnONPs against both H37Rv *Mtb* strain and MDR *Mtb* isolates, by blocking generation of crucial SufB protein via intein splicing inhibition.

A series of *in vitro* experiments, encompassing Bradford assays, UV-Visible spectroscopy, DLS, zeta potential, and Transmission Electron Microscopy (TEM), have collectively substantiated the interaction between the ZnONPs and the SufB protein. The Alamar Blue assay and Scanning Electron Microscopy (SEM) analysis further elucidated the bactericidal effects of the ZnONPs on H37Rv and MDR *Mtb* cells, which was correlated to the inhibitory effects of the ZnONPs on splicing and cleavage reactions of SufB precursor protein. Thus ZnONPs and the SufB protein interaction induced a significant alteration in the *Mtb* viability possibly due to loss of protein’s biological activity. While SEM confirmed extensive mycobacterial membrane damage, the Alamar blue assay highlighted the superior efficacy of a combination therapy (ZnONPs-INH/RIF & ZnONPs-LFX/MXF), where the effective dosage of the anti-TB drugs were reduced by half towards both drug-sensitive strain and MDR *Mtb* isolates ^82–85^. Next, *in vivo* experiments in infected mice model, validated substantial anti-mycobacterial activity of ZnONPs-RIF combination against the *Mtb* H37Ra strain.

In the current study, histopathological examination of infected lungs and spleen tissues (H&E staining) has provided evidence for the anti-inflammatory efficacy of ZnONPs (Figure 7). These results provided insight into the extent of inflammation and immune cell recruitment, which are the key indicators of the host immune response during *Mtb* infection and management ^103–105^. When administered individually, ZnONPs (26 µg/ml), ZnONPs (50 µg/ml), and RIF (15 mg/kg) caused a partial reduction in inflammatory cells within the infected lungs & spleen. On the contrary, mice treated with a combination of ZnONPs (26 µg/ml) + RIF (10 mg/kg) and ZnONPs (50 µg/ml) + RIF (10 mg/kg), demonstrated no inflammatory cell infiltrations, suggesting their superior anti-inflammatory action (Figure 7). The observed reduction in inflammatory cells in the ZnONPs + RIF treated mice group suggests a favourable anti-inflammatory and anti-mycobacterial effect ^106,107^.

These collective findings underscore the potential of ZnONPs as a promising additive along with existing anti-TB drugs in the fight against tuberculosis. Herein, we propose a dual mode of action for this ZnONPs-based combinatorial approach: ZnONPs can significantly influence mycobacterial viability by complete blockage of splicing and cleavage reactions to inhibit generation of essential protein SufB, further supported by the potent antimycobacterial action of anti-TB drugs (Figure 8). Since eukaryotes such as humans lack intein sequences, and microbial intein sequences resist mutations, targeted drug development strategy aiming splicing regulation of critical proteins have been proposed as novel mechanisms to manage drug-resistant infections ^108^. Current study also highlights the reduction in the effective dosage for the anti-TB drugs in combination with ZnONPs, that will address caveats such as drug toxicity. However, further research and clinical trials will be essential to validate the safety and efficacy of ZnONPs as a potential supportive treatment option for the infected patients.

**Figure 8.**
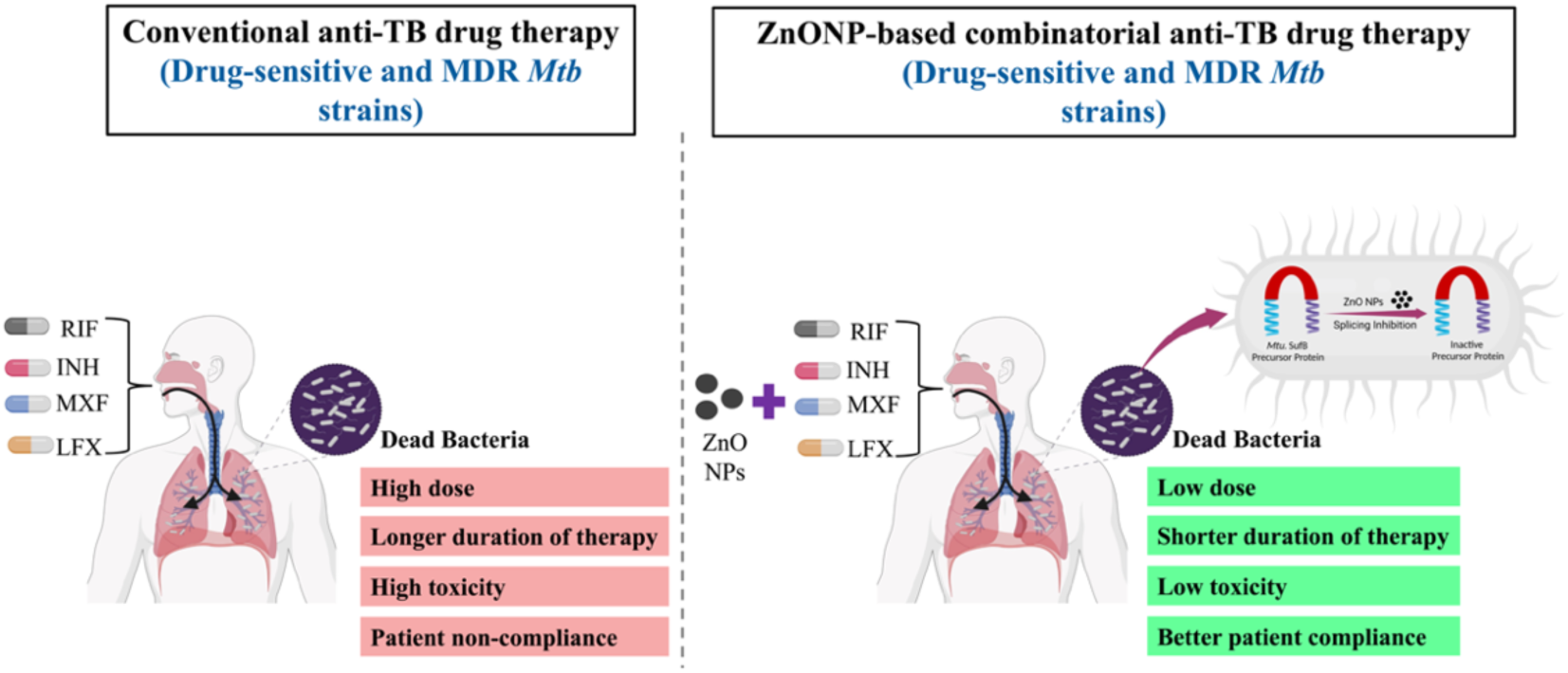
Schematic diagram illustrating the proposed mechanism for higher efficacy of combination therapy in managing drug-susceptible and MDR TB, contrasting with conventional drug therapy. ZnONPs and RIF/INH or MXF/LFX combination exhibits superior therapeutic potential possibly via a dual approach: targeting the mycobacterial viability by splicing regulation to inhibit generation of essential protein SufB, further supported by the potent antimycobacterial action of anti-TB drugs. Multidrug-resistant (MDR), Rifampicin (RIF), Isoniazid (INH), Moxifloxacin (MXF), Levofloxacin (LFX).

This study’s uniqueness lies in its innovative approach that demonstrated bactericidal efficacy against both drug-sensitive and MDR *Mtb* isolates, as a significant improvement over earlier research. Furthermore, the inhibition of SufB splicing by ZnONPs provides a new mechanism for managing TB, distinct from existing treatments. Additionally, the reduced dose of RIF/INH and MXF/LFX when combined with ZnONPs may lead to fewer adverse effects and improved patient compliance, especially addressing concerns during extended treatment periods (as in case of disseminated TB). Moreover, the potential toxicity of ZnONPs can be further eliminated by obtaining NPs via green synthesis routes ^109,110^. Howbeit, future research should focus on clinical trials to validate these findings and explore the long-term benefits and safety of this combinatorial approach. Current study addresses the growing issue of drug-resistant TB by exploring a new therapeutic approach that can have a significant contribution to the field of TB research and management. Higher efficacy along with lower drug dosage and probable shorter treatment duration may have a positive impact on the overall health cost, facilitating global effort to fight against TB.

## Author contributions

Deepak Kumar Ojha: Conceptualization, Resources, Data curation, Formal Analysis, Validation, Visualization, Methodology, Writing—original draft, Writing—review & editing; Ashwaria Mehra: Resources, Data curation, Funding acquisition, Validation, Methodology, Writing—review & editing; Sunil Swick Rout: Resources, Data curation, Software, Validation, Visualization; Sidhartha Giri: Resources, Validation, review & editing; Sasmita Nayak: Conceptualization, Resources, Data curation, Formal Analysis, Supervision, Funding acquisition, Validation, Investigation, Visualization, Methodology, Writing—original draft, Project administration, Writing—review & editing.

## Conflicts of interest

There are no conflicts to declare

## Data Availability

The data underlying the present article are available in the article and in its online Supplementary Information.

## Acknowledgements

The plasmids used in the present study are borrowed from Prof. Marlene Belfort’s Lab, SUNY, Albany, NY, U.S.A. Our sincere gratitude to the Central Instrumentation facility (CIF) at OUAT (Odisha University of Agriculture & Technology), Bhubaneswar and Dr. S.K Dash for his help in conducting SEM analysis at OUAT Bhubaneswar. We thank to Dr. Sravan Kumar for his help in conducting TEM analysis at the Central Institute of Petrochemicals Engineering and Technology (CIPET), Bhubaneswar. We extend our gratitude to Dr. Sidhartha Giri for providing access to the Biosafety Level 3 (BSL3) laboratory facility at ICMR-RMRC, Bhubaneswar, for smooth conductance of mycobacterial experiments.

## Ethics Statement

The animal study was reviewed and approved/recommended by the IAEC of Kalinga Institute of Industrial technology Deemed to be University (KIIT-DU), Bhubaneswar, Odisha. (Approval No. KSBT/IAEC/2023/2/14.2)

## Funding

This work is supported by Indian Council of Medical Research (ICMR), India (ICMR Project ID: IIRP-2023-7834/F1). A.M. is supported by INSPIRE fellowship (DST/INSPIRE/03/2021/002800/IF 190921); INSPIRE Division, DST, Government of India.

## Abbreviations

Ab, antibody; ANOVA, Analysis of Variance; CC, C-terminal cleavage product; IB, inclusion bodies; IPTG, isopropyl β-d-1-thiogalactopyranoside; I, intein; CC, C-terminal cleavage; P, precursor; LE, ligated extein; MDR, multidrug resistant; DS, Drug sensitive ; DR, Dug resistance; *Mtb*, *Mycobacterium tuberculosis*; NC, N-terminal cleavage product; NE, N-extein; Ni-NTA, nickel-nitrilotriacetic acid; PAGE, polyacrylamide gel electrophoresis; PVDF, polyvinylidene fluoride; SI, splicing inactive; SDS, sodium dodecyl sulfate; SEM, Scanning Electron Microscope; SUF, Mobilization of Sulphur; TB, Tuberculosis; TCEP.HCl, tris 2-carboxyl ethyl phosphine. Hydrochloric acid; TNB, 2-nitro-5-thiobenzoic acid; UV–Vis, UV– visible spectroscopy; WT, wild-type; NP, Nanoparticle; ZnO, zinc oxide; H&E; Hematoxylin and eosin.

## Notes

### Competing Interest Statement

The authors have declared no competing interest.

### Summary of Updates

Additional experiments like DLS and Zeta Potential included in the supplementary information. Figures in the main manuscript has been modified for clear understanding. Write up changes in the main manuscript and supplementary information for clarity.

