## Supplementary Information for "Bactericidal activity of ZnO nanoparticles-anti TB drugs combination towards H37Rv strain and multidrug-resistant isolates of *Mycobacterium tuberculosis* via SufB splicing inhibition"

### Table of Contents


| | |
|---|---|
| **Figure S1**. MTT assay to determine the cytotoxicity of ZnONPs | S3 |
| **Figure S2.** Dynamic light scattering (DLS) and zeta potential analysis of ZnONPs-protein complex. | S4 |
| **Table S1.** Hydrodynamic diameter (nm) and zeta potential (mV) measurements. | S4 |
| **Figure S3.** Confirmation of splicing and N-terminal cleavage products of *Mtb* SufB precursor via western blot. | S5 |
| **Figure S4.** Effects of intermediate concentration range of ZnONPs on *Mtb* SufB precursor splicing and N-terminal cleavage reactions. | S5-S6 |
| **Figure S5.** Determination of Minimum bactericidal concentration (MBC) for ZnONPs via Spread plate method. | S6 |
| **Figure S6.** Alamar blue assay to show the effects of ZnONPs on the growth and viability of *Mycobacterium smegmatis (M. sm)*. | S7 |
| **Reference** | S7 |

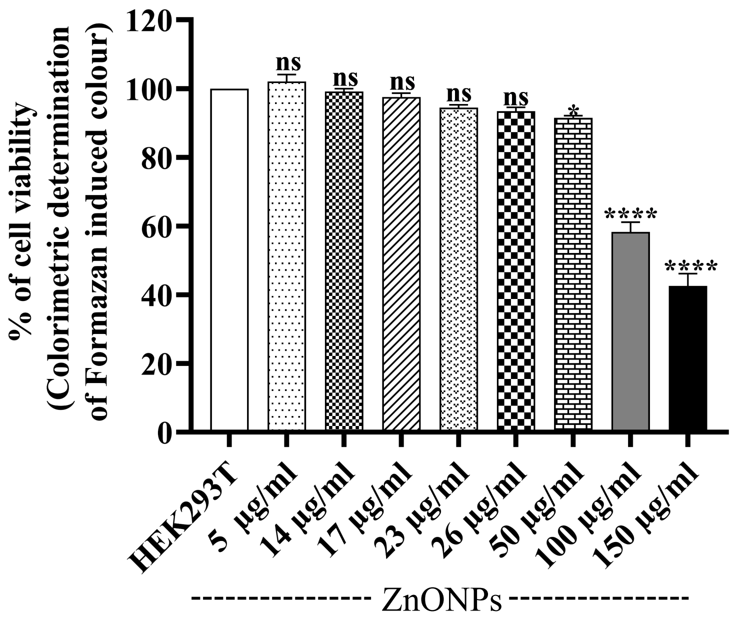


**Figure S1**. **MTT assay to determine the cytotoxicity of ZnONPs.** HEK293T cells (human embryonic kidney cell line) were exposed to varied concentrations of ZnONPs for 72 hrs, and the cell viability was determined as explained in the main text. Error bars represent (±1) SEM from (n=3) three independent sets of experiments.

**Dynamic light scattering (DLS) and zeta potential analysis of ZnONPs-protein complex**

Purified *Mtb* SufB precursor protein was refolded in the presence of different concentrations of ZnONPs (26 µg/ml and 50 µg/ml) as explained in section 2.7 for the *in vitro* refolding assay. Following incubation, the treated samples were analyzed via DLS and zeta potential (Malvern Nano – ZS90). To account for potential buffer-nanoparticle interactions, control samples included refolding buffer alone and the buffer with 50 µg/ml of nanoparticles were incubated under the same experimental conditions. The detailed analysis is provided in the Results and Discussion of the main text.

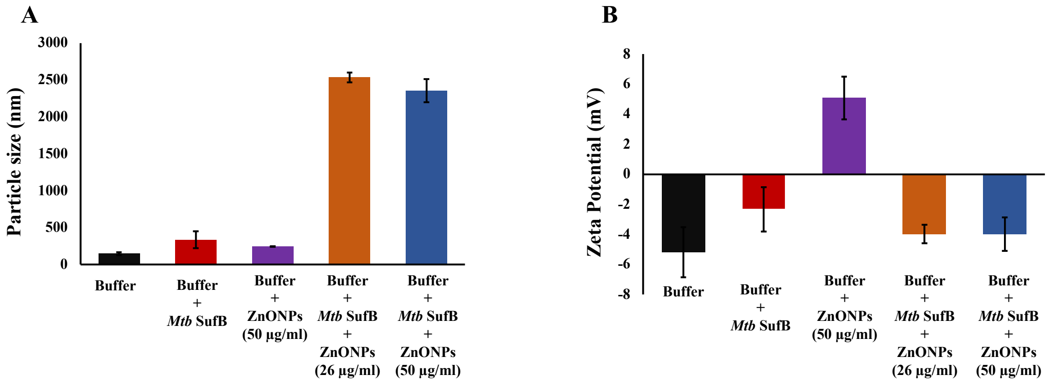


**Figure S2. Dynamic light scattering (DLS) and zeta potential analysis of ZnONPs-protein complex. (A)** Hydrodynamic diameter distribution. **(B)** Zeta potential profile. Error bars indicate ±1 SD for mean from three independent experiments (n=3).

**Table S1.** Hydrodynamic diameter (nm) and zeta potential (mV) measurements. Data represent mean (±1 SD) calculated from three independent experiments (n=3).

| Samples | Hydrodynamic diameter (nm) | Zeta Potential (mV) |
|---|---|---|
| Buffer | 144.9 ± 21.3 | -5.2 ± 1.7 |
| Buffer + *Mtb* SufB | 334.6 ± 115 | -2.3 ± 1.5 |
| Buffer + ZnONPs (50 µg/ml) | 242.7 ± 2.6 | 5.1 ± 1.4 |
| Buffer + *Mtb* SufB + ZnONPs (26 µg/ml) | 2535 ± 66.5 | -4.0 ± 0.6 |
| Buffer + *Mtb* SufB + ZnONPs (50 µg/ml) | 2355.3 ± 157.6 | -4.0 ± 1.1 |

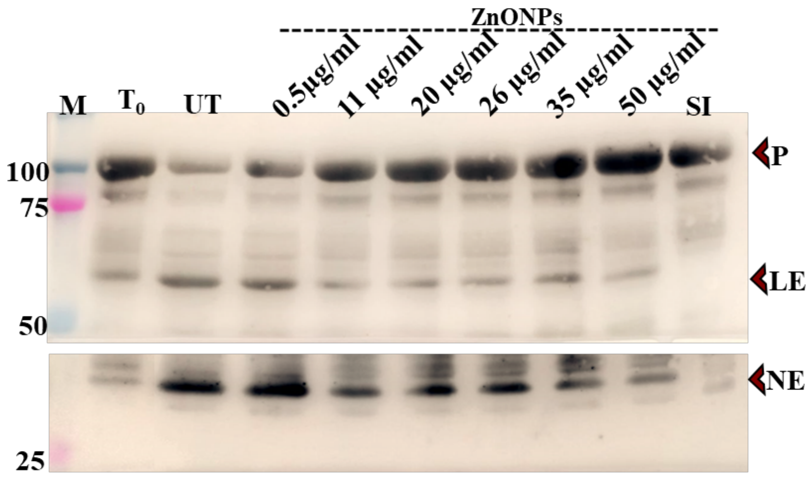


**Figure S3. Confirmation of splicing and N-terminal cleavage products of *Mtb* SufB precursor via western blot.** *Mtb* SufB precursor protein was refolded *in vitro* in presence varied concentrations of ZnONPs, resolved through 5%-10% gradient SDS PAGE, and the protein bands were detected by using anti-(His) antibody. The details of the methods and the results are mentioned in the main text. P: *Mtb* SufB precursor; LE: Ligated exteins; NE: N-extein; $T_0$: protein products at time 0, UT: untreated SufB precursor; M: protein molecular weight ladder

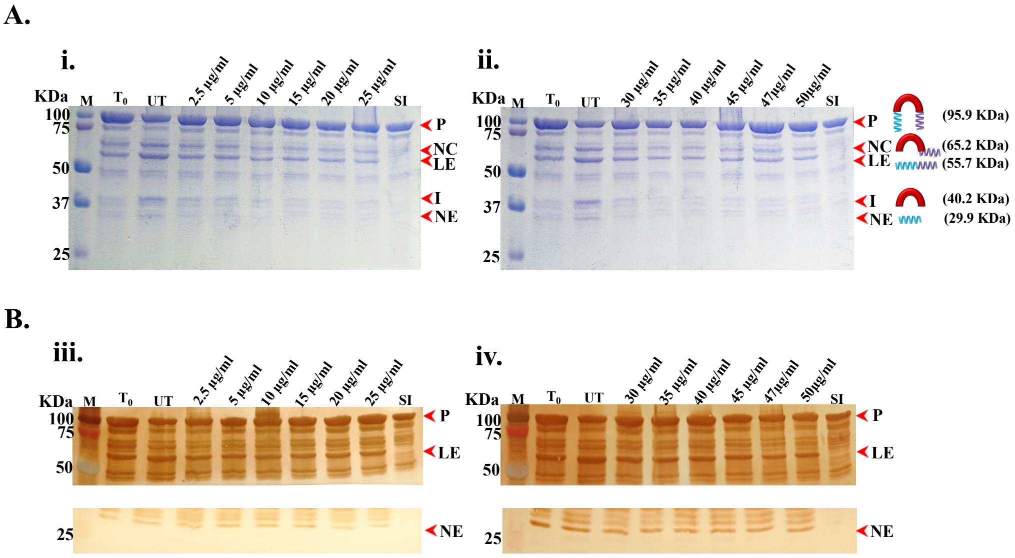


**Figure S4. Effects of intermediate concentration range of ZnONPs on *Mtb* SufB precursor splicing and N-terminal cleavage reactions. (A)** Products from *in vitro* protein refolding experiment were resolved through 4–10% gradient SDS-PAGE. ($T_0$): splicing and cleavage reactions at 0 h, (UT): untreated protein sample; and Lanes 4-9 show protein products induced by varied concentrations ZnONPs: (i) (2.5 µg/ml -25 µg/ml) (ii) (30 µg/ml to 50 µg/ml), Lane 10 (SI) for (i) and (ii): Splicing inactive SufB double mutant (Cys1Ala/Asn359Ala) is used as a negative control for splicing. **(B)** Western Blot analysis to validate the identity of protein products by using anti-(His) antibody. The details of the methods and the results are mentioned in the main text. P: *Mtb* SufB precursor; LE: Ligated exteins; NE: N-extein; $T_0$: protein products at time 0, UT: untreated SufB precursor; M: protein molecular weight ladder.

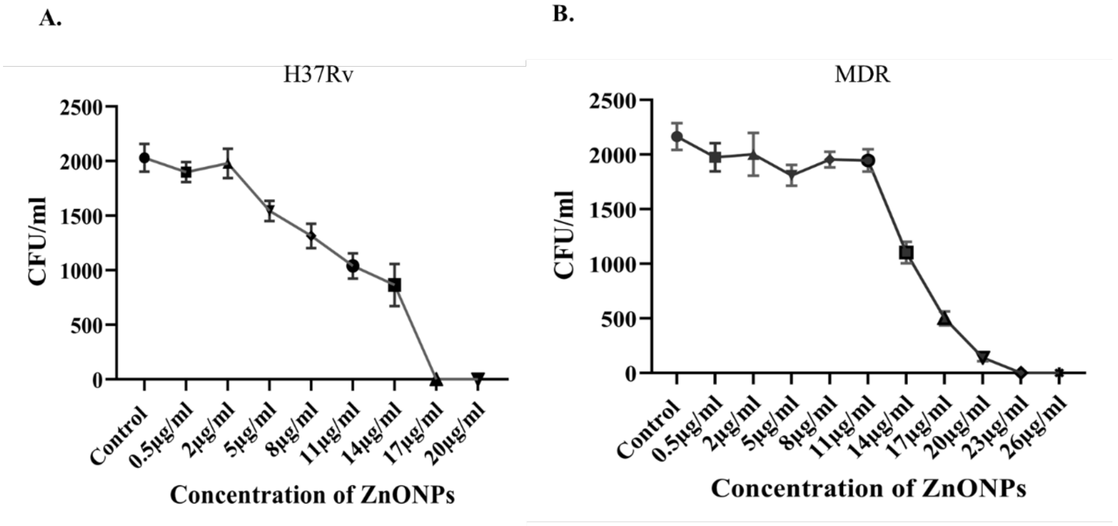


**Figure S5. Determination of Minimum bactericidal concentration (MBC) for ZnONPs via Spread plate method. A.** MBC of ZnO nanoparticles (ZnONPs) against H37Rv *Mycobacterium tuberculosis* (*Mtb*). **B.** MBC of ZnO nanoparticles (ZnONPs) towards multidrug-resistant (MDR) *Mtb isolates*. The details of the methods and the results are mentioned in the main text. Error bars represent (±1) SEM from (n=3) three independent sets of experiments.

#### Alamar blue assay to show the effects of ZnONPs on the growth and viability of *Mycobacterium smegmatis (M. sm)*

*Mycobacterium smegmatis* (*M. sm*) SufB precursor protein lacks an intein sequence in its polypeptide chain (1). Hence *M. sm* was used as a negative control to evaluate ZnONPs effect on its growth and viability, when intein splicing event is missing. All the cells exhibited pink colour in presence of ZnONPs, suggesting no effect of ZnONPs on *M. sm* viability due to lack of regulatory influence on SufB precursor splicing and cleavage reactions.

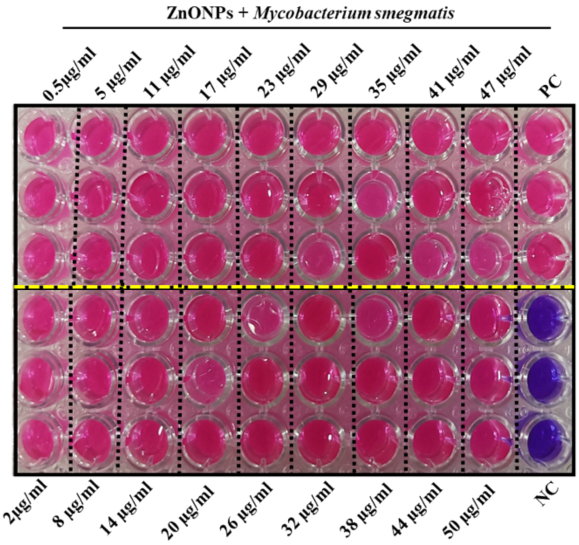


**Figure S6. Alamar blue assay to show the effects of ZnONPs on the growth and viability of *Mycobacterium smegmatis (M. sm)*.** $10^6$ CFU of bacteria were incubated with different concentrations of ZnONPs (0.5 µg/ml to 50 ug/ml). After 24h of incubation growth and viability of mycobacterial cells were analyzed by a colour change. The details of the methods and the results are mentioned in the main text. *M. sm* cells remained viable (pink) in presence of varied concentrations of ZnONPs. NC- Negative control (*M. sm* cells without ZnONPs), PC – positive control (*M. sm* cells + RIF 3 µg/ml)
